# Localgini: A method for harnessing inequality in gene expression to improve the quality of context-specific models

**DOI:** 10.1101/2023.09.14.557840

**Authors:** S Pavan Kumar, Nirav Pravinbhai Bhatt

## Abstract

Genome-scale metabolic models (GEMs) are valuable tools for investigating normal and disease phenotypes of biological systems through the prediction of fluxes in biochemical reactions. However, in specific contexts such as different cell lines, tissues, or diseases, only a subset of reactions is active. To address this, several model extraction methods (MeMs) have been developed to filter the reactions in GEMs and extract context-specific models. These methods utilize gene expression data as a source of context-specific information. To construct context-specific models, MeMs require core reactions specific to the given context as input. Typically, core reactions are derived using a single threshold applied to gene expression data. Reactions associated with genes whose expression values exceed the threshold are considered as core reactions. However, it is important to note that enzyme activity is not solely determined by gene expression levels. This approach based on a single threshold may inadvertently exclude reactions that require enzymes in smaller quantities. In this study, we propose a novel thresholding algorithm called *‘Localgini’*, which leverages the Gini coefficient and transcriptomics data to derive gene-specific thresholds. Localgini is implemented as a pre-processing step to obtain core reactions for MeMs. To demonstrate the effectiveness of Localgini, we construct context-specific models for NCI-60 cancer cell lines and human tissues using different MeMs. We compare the performance of Localgini with existing thresholding methods, namely LocalT2 and StanDep. The results show that the models derived using Localgini recover a greater number of housekeeping functionalities compared to the other thresholding methods. Moreover, the Localgini-derived core reactions exhibit increased self-consistency and display enhanced consensus among models built using different MeMs. By incorporating transcriptomic support, Localgini includes low-expression reactions in the core reaction list, enhancing the comprehensiveness of the resulting models. Codes used in this study, compatible with COBRA toolbox are available at https://github.com/NiravBhattLab/Localgini

**Author summary:** Genome-scale models are becoming a desirable tool to understand the metabolism of a biological system and hence find applications in the fields of systems and synthetic biology. These models are often integrated with transcriptomics data to improve prediction accuracy. Algorithms developed to integrate transcriptomics data with genome-scale models require core reactions to be derived from omics data using a threshold. In this work, we propose a thresholding method that uses an inequality-based metric to derive thresholds. We implied the proposed method and other existing methods to datasets of cancer cell lines and human tissue. We showed that our method improves the inclusion of reactions required for basic cellular maintenance. Furthermore, we validated the built models for the reduction in variance owing to the model-extraction algorithms. Overall, the proposed method improves the quality of metabolic models by inferring inequality in the distribution of gene expression levels across samples/contexts.

## Introduction

Metabolic networks play a crucial role in understanding biological systems, both in normal and disease conditions, by capturing the enzyme-catalyzed reactions, spontaneous reactions, and transport reactions under different environmental settings. To investigate these networks at a system-level, genome-scale metabolic models (GEMs) have been developed. These models mathematically represent the metabolic reactions in a biological system by formulating mass balance equations of metabolites, assuming a pseudo-steady-state condition [1]. By simulating GEMs, reaction fluxes under various physiological constraints can be estimated allowing to model of biological systems. In contrast, ordinary differential equations (ODE)-based models relax the steady-state assumption. However, they require defined kinetic parameters and initial metabolic concentrations for simulation studies and are computationally intensive for large-scale metabolic reaction studies [2]. Thus, the applications of ODE-based models for genome-scale studies are limited. Since GEMs do not require information on reaction kinetics, they are more suitable for studying large-scale metabolic systems such as multi-compartment cell systems [3], multi-tissue models [4], microbial communities [5], and disease conditions [6], albeit relying on the steady-state assumption [7].

In GEMs, the primary variable of interest is the fluxes through different metabolic reactions. GEMs have been in different applications such as strain design in biotech industries [8–10] drug target prediction in biomedical sectors as [11, 12] and investigating the molecular basis of diseases [13–15]. The study of human metabolic networks has significantly advanced with the development of GEMs such as Recon-X [16] and iHSA [17]. With the growing availability of omics data [18–21], the predictive capability of GEMs can be further enhanced using Model Extraction Methods (MeMs) developed in recent years [22]. MeMs integrate expression data specific to a particular context (e.g., cell type, disease state, or any perturbation affecting gene expression) and gene-protein-reaction (GPR) association rules to extract context-specific models. These methods provide a mechanistic link between gene expression data and phenotype of the given context. Context-specific models can be constructed either by constraining reactions in GEMs [23] (continuous methods) or by filtering out irrelevant reactions from GEMs (discrete methods). Discrete methods have advantages over continuous methods as they are robust to expression data noise and do not require data normalization [22]. Discrete methods involve discretizing gene expression data into core and non-core genes. Typically, a single threshold is applied to transcriptomics data to determine the core genes. Genes expressed above the threshold are considered the core genes, while those with expression values below the threshold are classified as the non-core genes [24]. MeMs extract a subset of reactions from GEMs, considering support for the given core reactions obtained using the core genes, objective function, stoichiometry matrix, and evidence from the existing literature.

Previous studies have highlighted that the choice of thresholding parameters significantly affects the final extracted model content [24, 25]. However, it is common practice to use a single threshold for all genes without much guidance in selecting appropriate threshold values. This approach can lead to the exclusion of housekeeping genes essential for maintaining cellular functionalities [24] when using a stringent threshold, or the inclusion of false positives when using a lenient threshold.

Additionally, using a single threshold overlooks variations in catalytic efficiencies of enzymes and complex pathway-specific expression patterns. To address these limitations, novel approaches are required to derive gene-specific thresholds for accurate context-specific metabolic modeling. If multiple samples are available, gene expression values across the samples can be utilized to compute gene-specific thresholds. One such approach is LocalT2, which calculates the mean of expression values as thresholds [26]. However, LocalT2 fails to capture the majority of housekeeping reactions in the core reaction list [27]. Another thresholding algorithm called StanDep [27] has been developed by Joshi et al., which hierarchically clusters genes based on their expression patterns and provides cluster-specific thresholds. The StanDep algorithm exhibits improved recovery of housekeeping functionalities and reduces false negatives in the core reaction sets. Nevertheless, determining appropriate values for hyperparameters such as bin edges, number of clusters, distance, and linkage metrics for hierarchical clustering in the StanDep algorithm remains a challenging task.

In this study, we propose a novel thresholding algorithm called *Localgini* which utilizes the concept of the Gini coefficient [28] to derive gene-specific thresholds. The outputs of the Localgini algorithm are further processed to obtain inputs for various discrete MeMs. We build context-specific models using Localgini in combination with six MeMs: FASTCORE [29], MBA [30], iMAT [31], INIT [32], GIMME [33], and mCADRE [34]. The performance of Localgini is compared to two gene-specific thresholding algorithms, LocalT2 and StanDep, using gene-expression datasets of cancer cell-line and Human Proteome Atlas (HPA). Our results demonstrate that the Localgini algorithm effectively incorporates a greater number of housekeeping functionalities into the context-specific models compared to other data-driven thresholding methods reported in the literature. Furthermore, the core-reaction list derived using Localgini exhibits higher self-consistency compared to other thresholding methods.

## Results

The Localgini method is applied to RNAseq data from NCI60 cancer cell lines [35] and the Human Protein Atlas [19] to obtain the core reaction set or reaction importance scores. These reaction importance scores are then utilized to extract context-specific models for a total of 44 cancer cell lines and 54 human tissues using various MeMs that have been previously published. The main results presented in the text primarily focus on the cancer cell-line data, while the findings related to human tissues are provided in the supplementary text (S1 Text).

### Localgini recovered more number of housekeeping reactions compared to other methods

Housekeeping genes are expressed uniformly across all cells, regardless of the specific context. These genes play a crucial role in maintaining cellular functions and are expected to have consistent expression levels across different conditions. In our study, a total of 929 housekeeping reactions are obtained by mapping housekeeping genes [36] to reactions using GPR rules in Recon2.2. To evaluate the ability of different thresholding methods to recover these housekeeping reactions, we compared the lists of core reactions derived from various thresholding approaches. Our analysis revealed that Localgini outperformed other methods in retrieving the highest number of housekeeping reactions in both the cancer cell-line data and the Human Protein Atlas (HPA) data. Among the 44 cancer cell lines analyzed, Localgini successfully retrieved the most housekeeping reactions in 35 of them, as depicted in Figure 2. Similarly, among the 54 human tissues studied, Localgini retrieved the most housekeeping reactions in 31 tissues (refer to S3 Fig). Since housekeeping genes exhibit constant expression values across different samples, the corresponding reactions are not specific to any particular context.

**Fig 1.**
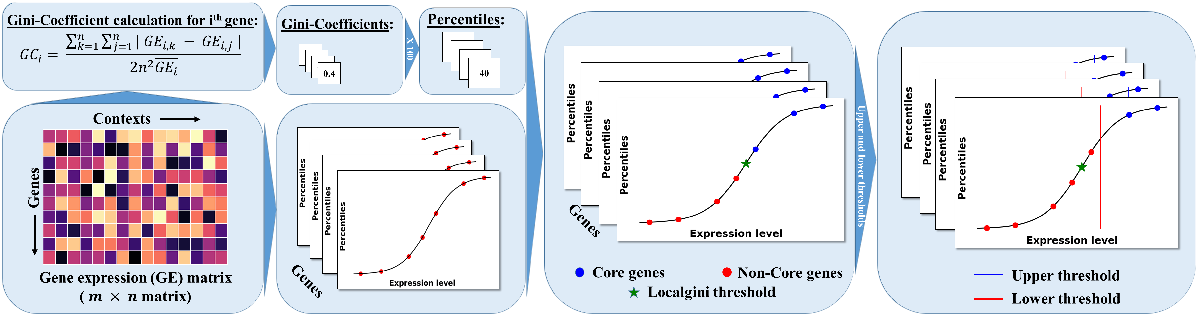
Implementation of Localgini thresholding method on a transcriptomic data to define core/non-core reactions.

**Fig 2.**
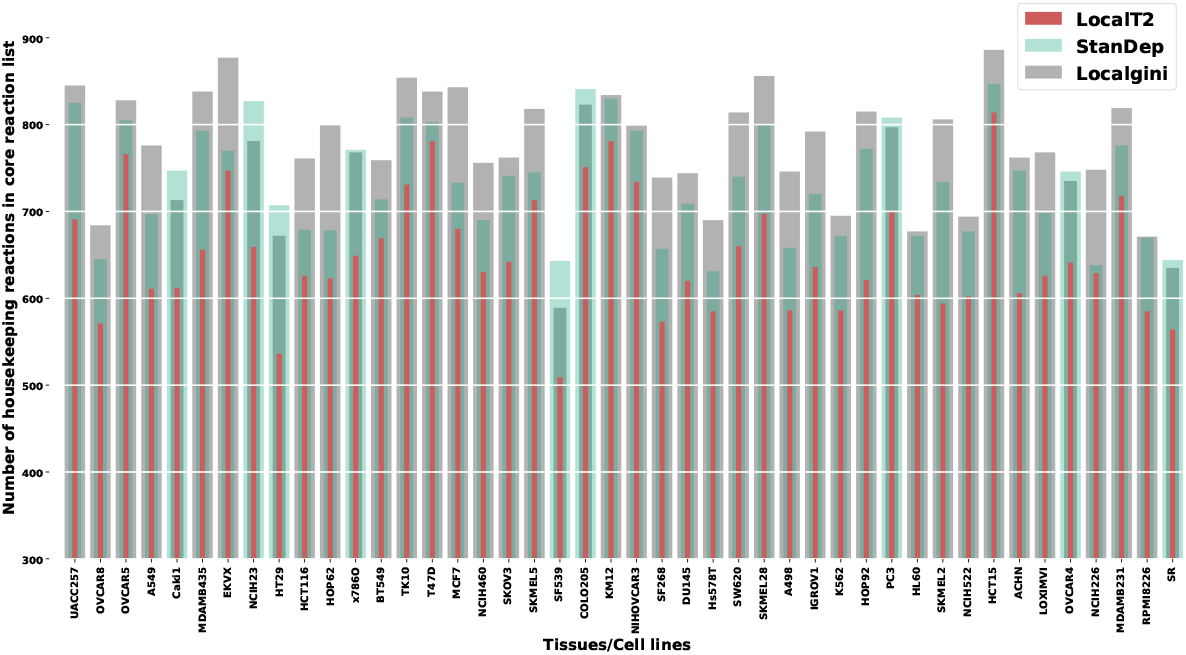
Number of housekeeping reactions rectified in core reaction list by each of the thresholding methods in NCI60 cancel cell-lines data.

Consequently, the threshold for these reactions should be set at the minimum level required to support their activity across all contexts. The Gini coefficients associated with housekeeping reactions are expected to be low due to the equal distribution of expression levels across different contexts. Given that Localgini is designed based on the concept of the Gini coefficient, it effectively captures these context-non-specific reactions across diverse cell lines. Therefore, Localgini includes a significant portion of these housekeeping reactions in different cell lines, irrespective of the specific context.

### Localgini derived core reaction list are self-consistent

A reaction is included in context-specific models from GEMs due to three factors: (i) constraints on exchange reactions, (ii) a predefined core reaction list, and (iii) the inclusion of the reaction by the MeM, which supports the core reaction list. Recently Joshi et al. [27] demonstrated that the MeM has a significant impact on the final extracted model content compared to the constraints. Therefore, a reaction included in the final context-specific model must originate from either the predefined core reaction list or the applied MeM. In the presence of a core reaction list, the MeM attempts to add or remove specific reactions from it to obtain the final model following different assumptions. Self-consistency refers to a property of the thresholding method used to assess the fraction of reactions in the final model that comes from the thresholding process. 1764 models were constructed for 44 cancer-cell lines and 54 human tissues by employing three threshold types and six model extraction methods. Figure 3 and S4 Fig illustrate that core reaction lists derived from Localgini exhibit higher self-consistency in most of the MeMs, with a majority of reactions in the model stemming from the thresholding method, compared to other thresholding methods. The distribution of the fractional contribution of reactions from MeMs using Localgini is compared with the distributions obtained using LocalT2 and StanDep. To compare the distributions obtained using Localgini and LocalT2 and those obtained using Localgini and StanDep, a left-tailed Wilcoxon rank-sum test is conducted. The distribution obtained using Localgini is significantly lower (*p*-value*<* 0.01) than the distributions obtained using LocalT2 and StanDep in all MeMs, except for mCADRE in the case of NCI60 cancer cell-line data (Figure 3). For the human tissue dataset, the distribution obtained using Localgini is comparable to the distribution obtained using StanDep in FASTCORE and mCADRE. Additionally, the distribution obtained using Localgini is similar to the one obtained using LocalT2 in GIMME. LocalT2 yields more self-consistent reactions in iMAT in comparison to the other two methods (refer to S4 Fig).

**Fig 3.**
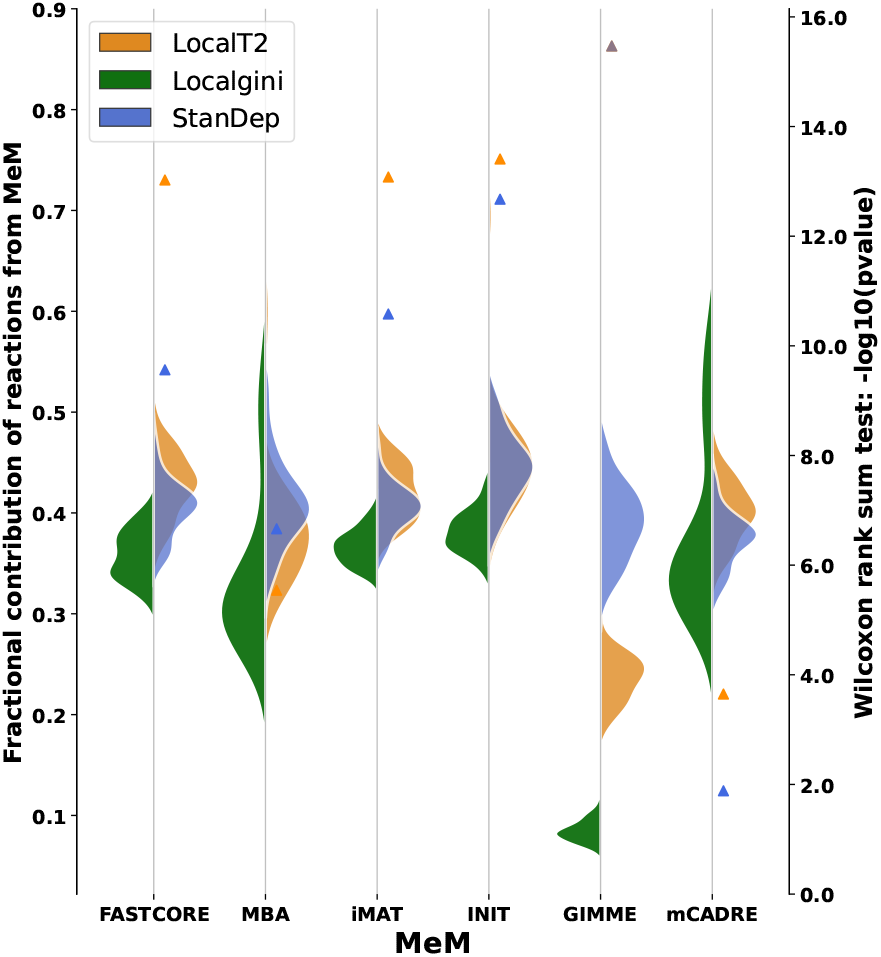
Localgini derived models are more self-consistent compared to other thresholding methods. Left y-axis: Violin plots indicating the fraction of reactions added by MeMs for the given reaction importance by different thresholding methods. Right y-axis: Triangles indicating the −*log*_10_(*p*−*value*) of the left-tailed Wilcoxon rank sum test to compare the distributions of Localgini with those of LocalT2 (orange colour) and StanDep (blue colour)

### Gene essentiality predictions for NCI60 cancer cell-lines

To identify essential genes in the context-specific models, we employed the Fast-SL algorithm [37]. The Fast-SL algorithm narrows down the search space of genes and iteratively predicts the genes necessary for cell growth. A gene is considered essential if the ratio of the maximum biomass flux after removing the gene to the maximum biomass flux before removing the gene is less than 0.05. To validate the predicted essential genes, we compared them with CRISPR-Cas9 loss-of-function screens data [38, 39]. In this dataset, each gene is assigned a score that quantifies the impact on cell growth when the corresponding gene is removed. A more negative score indicates higher importance of the gene for the growth of the respective cancer cell. To evaluate the performance of the predicted essential genes, we compared them with the negative CRISPR/Cas9 scores. As illustrated in S7 Fig, the Localgini method demonstrated comparable performance to the other methods in predicting gene essentiality. The Localgini method successfully identified essential genes in a manner similar to the other evaluated methods.

### Localgini reduces false negatives in the core reaction list

We performed pathway enrichment analysis on the core reaction lists obtained by applying all three thresholding methods to the Human Protein Atlas (HPA) dataset. The core reaction list was compared against known pathways associated with the respective tissues. A pathway is considered enriched in a specific tissue if the resulting *p*-value from the hypergeometric test is less than 0.05. The hypergeometric test was conducted using the *hygecdf* function in MATLAB, with inputs including the total number of reactions in Recon2.2 (*M*), the total number of reactions in Recon2.2 from the desired pathway (*K*), the number of reactions in the core reaction set (*N*), and the number of reactions in the core reaction set from the desired pathway (*x*).

The known pathways associated with the tissues were obtained from the supplementary data of [25], which included a total of 154 known pathway-tissue pairs. Among these pairs, there were 32 unique tissues and 29 unique pathways. Of the 54 human tissues in the HPA dataset, 31 tissues overlapped with the pathway-tissue pairs data.

Based on our analysis, the core reaction lists derived using the Localgini method exhibited enrichment with a greater number of pathways known to occur in the corresponding tissues. This finding is supported by Table 1 and S8 Fig in the supplementary information. The Localgini method demonstrated superior performance in capturing pathways that are known to be associated with specific tissues.

**Table 1.**
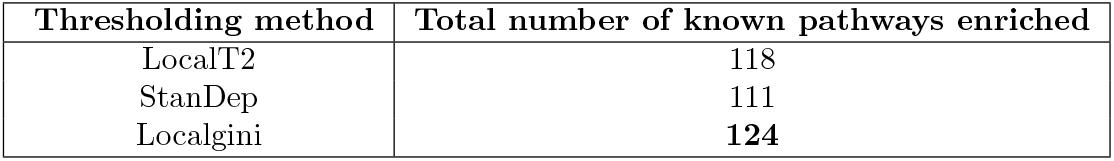
Localgini reduced false negatives compared to other thresholding methods.

| Thresholding method | Total number of known pathways enriched |
| --- | --- |
| LocalT2 | 118 |
| StanDep | 111 |
| Localgini | **124** |

### Localgini shows less variance across models built from the same cell-line data

In this study, different Model extraction Methods (MeMs) were employed, each utilizing distinct optimization algorithms based on underlying assumptions, to build context-specific models. These assumptions led to significant variations in the models generated from the same transcriptomics data or cell lines but using different MeMs. To address this variation and evaluate the impact of pre-processing decisions, such as the choice of thresholding method and the derivation of inputs to MeMs, on model consensus, two sets of matrices were utilized. The first set of matrices was used to explain the variance in reaction content among the different models, while the second set of matrices aimed to explain the variance in metabolic functionalities. Principal Component Analysis (PCA) was performed on these matrices to identify the principal components that contributed to the observed variances. The contribution of variance in each principal component was quantified for different categories, including MeM, cell lines, and cancer types. By conducting PCA and analyzing the contribution of variance in the principal components, the study aimed to assess the impact of different factors, such as the choice of MeM, specific cell lines, and cancer types, on the resulting models. This analysis provided insights into the extent of consensus achieved among the models and helped evaluate the effectiveness of various pre-processing decisions in building more consistent and reliable context-specific models.

### PCA on reaction content matrix

Variance from cancer-type and cell-line in PC1 is maximum in Localgini models. One more binary matrix is created where only the MeMs that try to include the core/active reactions in the final model (FASTCORE, MBA, and mCADRE) are considered. Other methods either try to support biomass objective (GIMME) or find a trade-off between high-confidence and low-confidence reactions (iMAT and INIT). In this matrix, the maximum contribution of variance in PC1 is from cancer-type in both Localgini (36.4%) and StanDep (28.4%). For LocalT2, the maximum contribution of variance in PC1 is from cell-line type (32.65%). Cell-line type contributed the second maximum variance in PC1 for Localgini (33.89%) and StanDep (26.51%) built models. Cancer type contributed the second maximum variance in PC1 for LocalT2 models (31.45%).

To analyze the variance in the models and understand the contributions of different factors, a binary matrix was constructed. This matrix had rows representing the models and columns representing the reactions in Recon2.2. The elements in the matrix indicated the presence (1) or absence (0) of a reaction in a particular model. To reduce dimensionality, PCA was performed on the matrix. Before conducting PCA, Columns with zero variance are removed and each row of the matrix was centered to have a zero mean. PCA allowed for the identification of the principal components that explained the most significant variances in the data. The analysis revealed that MeMs induced the maximum variance in the Principal Component 1 (PC1) of the reaction content matrix across all the thresholding methods (Figure 4). This indicated that MeMs played a crucial role in driving the differences among the models. Interestingly, variance from cancer-type and cell line in PC1 is maximum in Localgini models, suggesting that these factors had a significant influence on the resulting models. Another binary matrix was created, considering only the MeMs that aimed to include the core reactions in the final model, namely FASTCORE, MBA, and mCADRE. Other methods, such as GIMME, focused on supporting the biomass objective, while iMAT and INIT sought to find a balance between high-confidence and low-confidence reactions. In this new matrix, the maximum contribution of variance in PC1 was observed to come from cancer-type in both Localgini (36.4%) and StanDep (28.4%) models. For LocalT2 models, the maximum contribution of variance in PC1 came from the cell line type (32.65%). The second highest variance contribution in PC1 for Localgini (33.89%) and StanDep (26.51%) models came from the cell line type. In LocalT2 models, the second highest variance contribution in PC1 came from the cancer-type (31.45%).

**Fig 4.**
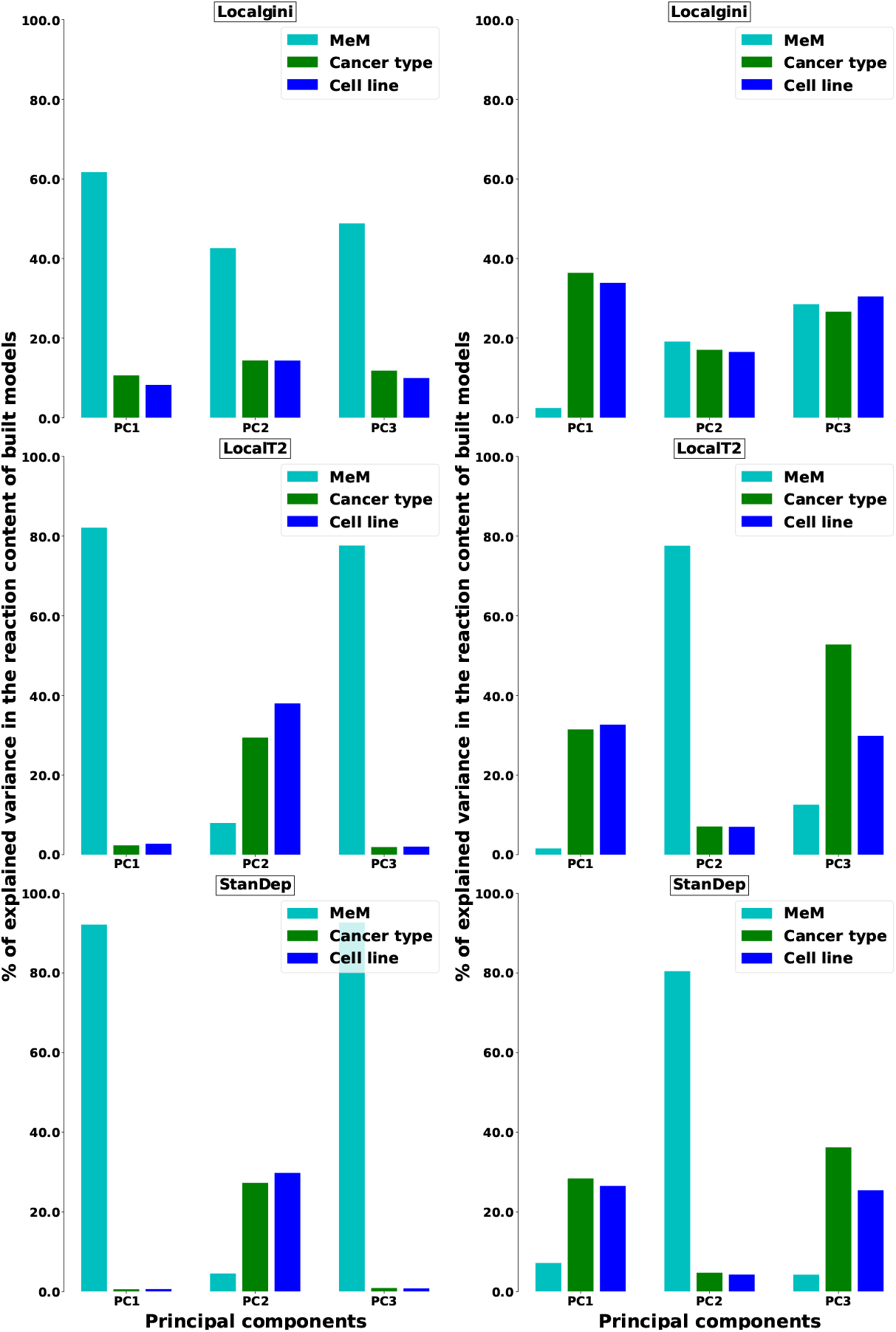
Models built from different MeMs show increased consensus in terms of reaction content in Localgini. Bar plots show the contribution of variance in the model’s reaction content from different categories considered. In the first column of plots, all MeMs are considered and in the second column of plots Only FASTCORE, MBA, and mCADRE are considered

### PCA on metabolic functionality matrix

To assess the variance in metabolic functionality of the models, a binary matrix was constructed, with rows representing the models and columns representing the 210 metabolic tasks curated by Richelle et al. [26]. The elements of the matrix indicated whether a particular model passed (1) or failed (0) the corresponding metabolic task. Similar to the analysis of the reaction content matrix, PCA was performed on this binary matrix to identify the main sources of variance among the models. The aim was to understand the contributions of different factors, such as MeMs, cancer-type, and cell line, to the variance in metabolic functionality. The analysis revealed that MeMs had a greater contribution to the variance in the models’ metabolic functionality compared to other categories. Furthermore, when examining the variance in Principal Component 1 (PC1), it was found that variance from cancer-type and cell-line is highest in Localgini compared to other methods in both cases of MeMs used (Figure 5).

**Fig 5.**
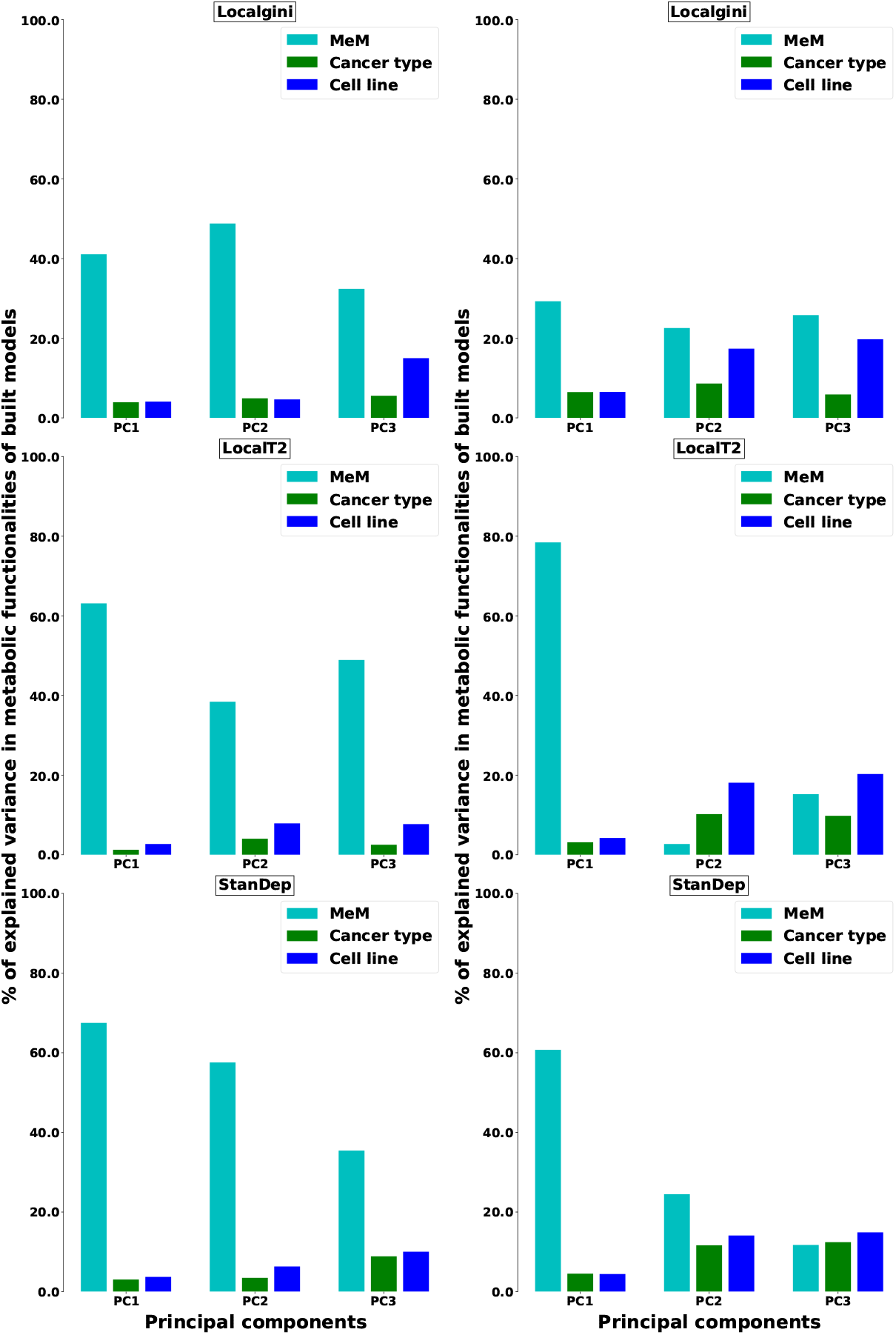
Models built from different MeMs show increased consensus in terms of metabolic functionalities in Localgini. Bar plots show the contribution of variance in the model’s metabolic functionality from all different categories considered. In the first column of plots, all MeMs are considered and in the second column of plots, Only FASTCORE, MBA, mCADRE are considered

## Discussion

Model extraction methods(MeMs) encompass a spectrum of algorithmic approaches employed alongside predefined reaction importance scores and GEMs to build context-specific models. The manual curation of reaction importance scores across the vast array of reactions inherent in Recon2.2 is undeniably laborious. Consequently, the prevailing practice involves harnessing omics data to define the active reaction set (or) reaction importance scores. The role of reaction in a particular context depends on gene expression level and different processes such as translation, environmental factors, and availability of substrates for the reaction. Hence, providing a single threshold to all the genes for defining the active reaction set is an abstraction that may inadvertently neglect reactions catalyzing stable proteins or those necessitating trace amounts. Should gene expression data span multiple contexts, a gene-specific threshold can be obtained by scrutinizing the gene’s variations across these contexts. The proposed approach, Localgini, is one such thresholding method that analyses the inequality in the distribution pattern of the gene expression across contexts to derive the gene-specific thresholds.

The Localgini-derived core reaction list captured the housekeeping functions better in comparison to other thresholding methods (refer to Figure 2 and S3 Fig). Moreover, we analyzed the core reaction set from cancer cell lines for housekeeping reactions captured only by Localgini (not other methods in any cell line). For this analysis, the housekeeping reactions captured by LocalT2 and StanDep in all the cell lines are removed from the Localgini-derived housekeeping reactions. The resulting reactions belong to amino acid metabolism, fatty acid oxidation, sphingolipid metabolism, and pyrimidine synthesis. Furthermore, models built from different thresholding methods are validated for housekeeping reactions recovery. Models extracted by combining Localgini and MeMs such as INIT and iMAT (that works by finding the trade-off between including core reactions and excluding non-core reactions) removed a few housekeeping reactions, from the core reaction set. All other models included a few more housekeeping reactions. Also, the models with Localgini as the thresholding method recovered the most number of housekeeping reactions in MeMs such as Fastcore, GIMME, and INIT compared to other thresholding methods (refer to S5 Fig and S6 Fig). We also checked whether the models are able to recover all the cancer hallmark genes [40]. Localgini-derived models are able to recover the most number of hallmark genes in all MeMs except MBA. In MBA, Localgini-derived models are comparable with StanDep-derived models in hall-mark gene recovery(refer to S14 Fig).

The reaction contribution of MeMs to the final model is less in the Localgini-based models (refer to Figure 3 and S4 Fig) compared to the LocalT2 and StandDep-based models. This has been verified using the Wilcoxon one-tailed rank-sum test. It can be interpreted as follows: Localgini provides expression support to the reactions that are not included in the core reaction list or otherwise included by the MeMs of other thresholding methods.

MeMs used in this study make different assumptions and use various algorithms to extract models. Hence, we investigated whether the Localgini-derived cancer cell-line models show less variability across the model extraction algorithms. Accordingly, variance explained by cancer-type and cell-line type is maximum in Localgini-derived models in PC1 compared to other methods (Figures 4 and 5). Moreover, the Jaccard similarity index for models extracted by different MeMs for each cell line is calculated for all the thresholding methods. The Localgini-derived models show increased consensus among them compared to other methods when all the MeMs were considered (refer to S10 Fig and S12 Fig). When only FASTCORE, MBA, and mCADRE are considered, all three thresholding algorithms have a similar distribution of Jaccard similarity scores (refer to S11 Fig) and S13 Fig)

Localgini and other thresholding methods are also implied on the HPA dataset and are evaluated for their ability to enrich the core reactions with well-studied pathways in the respective tissues. In the case of ubiquitous pathways (present in all the tissues), core reactions from all the thresholding methods are enriched in glycolysis/gluconeogenesis pathway reactions (refer to S8 Fig). However, reactions from the pentose phosphate pathway that generates pentose sugars, NADPH, and precursors for nucleotide synthesis are mostly enriched in the Localgini-derived core reaction list.

Hyperparameters for Localgini are lower and upper thresholds to exclude low-expression genes and include high-expression genes, respectively. These parameters are chosen based on the false negativity rate done in a previous study on LocalT2 using the same dataset [25, 26]. On the other hand, the hyperparameters in StanDep are comparatively high. These parameters are bin edges, distance metric, linkage metric for hierarchical clustering, and choice of a number of clusters, and the performance of the clustering method depends on the choice of hyperparameters.

The thresholding methods are evaluated based on the inclusion of context-relevant reactions. Localgini performs better with MeMs methods, Fastcore, GIMME, and iMAT, in comparison to Local T2 and StandDep Localgini gives a transcriptomic explanation of using inequality metric in threshold calculation. Localgini calculates a gene-specific threshold, and hence, it circumvents the oversimplified assumption on expression patterns of genes.

## Methods

### Data sets and software used

The study utilizes RNA-seq datasets from the NCI60 cancer cell lines, which consist of gene expression levels of 44 cancer cell lines representing 9 different cancer types [35]. In addition, the Human Protein Atlas (HPA) dataset is used, which includes consensus transcript expression levels summarized per gene across 54 tissues based on transcriptomics data from HPA and GTEx [19]. The proposed method, Localgini, is implemented and compared with the LocalT2 and StanDep approaches. These approaches involve different thresholding methods and are implemented using MATLAB 2022a. To validate the models built using different thresholding methods, several criteria are considered. The recovery of housekeeping reactions is assessed using housekeeping genes data from Eisenberg et al. [36]. The retrieval of cancer hallmark genes is evaluated using data from Liberzon et al. [40]. Furthermore, gene essentiality predictions are validated using CRISPR/Cas9 loss-of-function screens data from Aguirre et al. [39], Meyers et al. [38], and Doench et al. [41]. The genome-scale metabolic model (GEM) used in this study is Recon2.2, which represents human metabolism. Recon2.2 is downloaded from the Biomodels database [42]. The biomass production reaction in Recon2.2 is modified to include the metabolites required for growth, as described in a previous study. [24]. For the simulations, the COBRA toolbox 3.0 is employed, and the solver used is CPLEX.

### Data processing

To extract context-specific models using the Localgini method, the first step involves processing the gene expression data. Two alternative approaches can be employed: i) Thresholding gene expressions: In this approach, thresholds are applied to the gene expressions, followed by mapping them to reactions using gene-protein-reaction (GPR) rules. Once the core genes are obtained, they are mapped to the corresponding reactions based on the GPR rules provided in the Recon2.2 model. ii) Thresholding reaction expressions: In this approach, the gene expression values are first mapped to reactions using the GPR rules, and then thresholds are applied to the reaction expressions. Recently, [27] have used a different approach to providing thresholds for enzyme expressions by mapping only for the ‘AND’ relation in GPR rules. In the case of the Localgini method, the thresholding step is performed at the gene level, followed by mapping the core genes to reactions using the GPR rules provided in the Recon2.2 model (more details in supplementary text). For the GPR mapping, *min* of expression values is used for the AND rule, and *max* of expression values is used for the OR rule.

For the cancer cell-line models, specific considerations are made. Twenty-seven exchange reactions, each representing the secretion of a different metabolite, are included in the Recon2.2 model. The uptake and secretion bounds of seventy-eight exchange reactions are constrained based on exo-metabolomics data specific to the respective cancer cell lines [24, 26, 43]. The flux bounds of all other reactions in the model are set to −000 (lower bound) and 1000 (upper bound) for reversible reactions, and 0 (lower bound) and 1000 (upper bound) for irreversible reactions. To ensure flux consistency, a flux-consistent model of the Recon2.2 reconstruction is extracted for each of the 44 cancer cell lines using the *fastcc* [29] function, with a reaction activity cutoff set to 1×10^−8^. The obtained flux-consistent models of the cancer cell lines are then used as inputs to the MeMs.

For the human tissue models, the flux-consistent model of Recon2.2 is used as input for all the MeMs.

The biomass production reaction (‘Biomass reaction’) and ATP hydrolysis reaction (‘DM atp c’) are prevented from being removed in all the context-specific models. The lower bounds of biomass production and ATP hydrolysis reactions of the input Recon2.2 are set to 0.01 and 1.833, respectively. Additionally, housekeeping genes obtained from Eisenberg et al. [36] are mapped to reactions in Recon2.2 to identify the housekeeping reactions.

### Localgini: Gini coefficient based thresholding method

The Localgini method is a gene-specific thresholding method based on the concept of the Gini coefficient (GC) (Eq (1)). It calculates different threshold values for different genes based on their Gini coefficients. The Gini coefficient (∈ [0, 1]) measures the inequality in the dispersion of gene expression values across different contexts. A higher Gini coefficient indicates greater inequality, meaning that the gene is not uniformly expressed across all contexts. On the other hand, a lower Gini coefficient suggests a more equal dispersion of expression values, indicating a context-non-specific gene. To apply the Localgini method to a gene expression matrix (**GE**) of size *m* × *n* (where *m* is the number of genes and *n* is the number of contexts), the following steps are performed (refer to Figure 1):

1. Compute the Gini coefficient (*GC*) for each gene in the (**GE**) matrix using the formula.

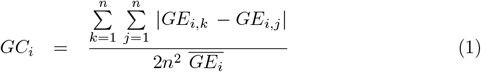

where *GC*_*i*_ represents the Gini coefficient of the *i*th gene, *GE*_*i,k*_ and *GE*_*i,j*_ are expression values of the *i*th gene for the *k*th and *j*th contexts, respectively, and 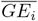 is the mean expression of the *i*th gene across contexts. The Gini coefficients are then stored in a vector **GC**.
2. Multiply the vector **GC** by 100 to obtain percentage values, resulting in the vector **GCP**.
3. Calculate the Localgini threshold (*LG*_*i*_) for each gene as the 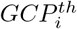 percentile value of *GE*_*i*_. This is done by sorting the expression values of the *i*th gene in increasing order and determining the 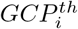 percentile using linear interpolation [44] as shown in the following example.
  - The gene expression values are sorted in increasing order and the *j*^*th*^ sample is taken as 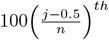 percentile value.
  - Linear interpolation is used to get the gene expression value for a given percentile. For example, if *GCP*_*i*_ = 15.5 for *i*^*th*^ gene with expression vector [91.51, 48.82, 57.06, 83.88, 49.46, 62.85, 40.95] then *LG*_*i*_ is calculated from 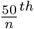 and 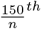 percentile values (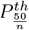 and 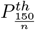).

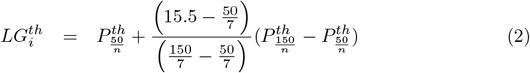

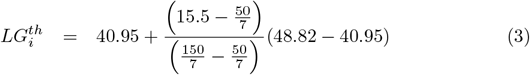

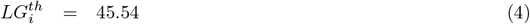
4. Two more thresholds upper (*U*) and lower (*L*) are given as global thresholds to the gene expression matrix, **GE**. The gene-specific threshold (*TH*_*i*_) for the *i*th gene is then determined based on the Localgini threshold (*LG*_*i*_) and the upper and lower thresholds using the following conditions:

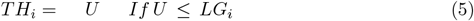

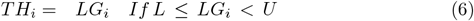

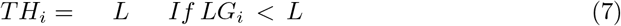
5. Identify the active/core gene set by considering the expression values of genes greater than *TH*_*i*_.

Including the upper and lower thresholds in the Localgini algorithm helps address potential limitations when processing genes with extremely low or high expression values. The upper and lower thresholds act as global thresholds that include high-expression genes (if greater than the upper threshold) or exclude low-expression genes (if lesser than the lower threshold) in the core gene set, irrespective of the gene-specific Localgini threshold. For this study, the 25^*th*^ percentile and 75^*th*^ percentile of the **GE** matrix is used as the lower and upper thresholds, respectively.

### Implementation of Localgini with model extraction methods

To extract context-specific models using the Localgini method, reaction importance needs to be provided in a specific format that is compatible with the algorithm used by the MeM. In this section, the implementation of Localgini to derive reaction importance for context-specific models is discussed. For the Localgini method, the gene-specific thresholds obtained from the Localgini algorithm are used to filter the reactions in the genome-scale metabolic models (GEMs). The reactions are considered “important” if their associated genes have expression values above the gene-specific threshold.

Reactions with expression values below the gene-specific threshold are considered “non-important” and are filtered out. In addition to the gene-specific thresholds, certain reactions are assigned higher importance in all the context-specific models. These reactions include the biomass function reaction, which represents the production of biomass components necessary for cell growth, and the ATP demand reaction, which represents the hydrolysis of ATP for energy production. These reactions are essential for cell survival and are typically assigned higher importance in all the models. The implementation details for providing reaction importance in the LocalT2 and StanDep methods can be found in the method section of supporting information. The detailed description can be found in the method sections of [26] for the LocalT2 method and [27] for the StanDep method.

### Core/non-core reaction sets

In the FASTCORE [29] and iMAT [31] methods, the concept of core and non-core reaction sets is used to build context-specific models. Here’s how these methods utilize the core and non-core reaction sets, specifically in relation to the Localgini thresholds:

In FASTCORE, all the reactions with expression values greater than the Localgini thresholds are considered as core reactions. These core reactions are considered essential and are included in the context-specific model. All other reactions that have expression values below the Localgini thresholds are considered non-core reactions. In iMAT, the determination of core and non-core reaction sets is slightly different. Reactions that have no details of gene-protein-reaction (GPR) associations in the Recon2.2 reconstruction and reactions with no available gene expression data (NER) are not included in either the core or non-core reaction sets. The reactions with expression values greater than the Localgini thresholds are considered as core reactions. All other reactions, which have both GPR associations and gene expression data, are included in the non-core reaction set of iMAT.

### Reaction importance scores

In the INIT [32], GIMME [33], and mCADRE [34] methods, reaction importance scores (*RIS*_*r,c*_) are calculated for every reaction in the genome-scale metabolic models (GEMs) based on the expression values and Localgini thresholds. The calculation of *RIS*_*r,c*_ is as follows:

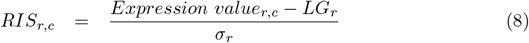

In this equation 8, *Expression value*_*r,c*_ represents the expression level of the *r*th reaction in the context *c, LG*_*r*_ is the reaction-specific Localgini threshold obtained from the gene-specific thresholding step, and *σ*_*r*_ is the standard deviation of the expression values of reaction *r* across all the contexts. The purpose of calculating the reaction importance scores is to quantify the relative importance of each reaction in a specific context based on its expression level compared to the Localgini threshold and the variability of its expression across different contexts. A positive RIS indicates that the reaction is highly expressed compared to the Localgini threshold, while a negative RIS indicates a lower expression. To handle extreme values, reactions with RIS less than −0 are assigned a fixed value of −0. This thresholding helps to limit the impact of outliers or very low expression values on the calculated RIS. RIS is further processed to get MeM-specific input (refer to S1 Text).

### High/Medium confidence reaction sets

In MBA [30] method, medium confidence reactions (*M*) and high confidence reactions (*H*) are defined based on the reaction importance scores (*RIS*_*r,c*_) calculated using the Localgini method. The process of defining these sets is as follows:

Reactions with positive RIS values: All reactions with positive RIS values are considered for inclusion in either the *M* or *H* set, depending on their percentile rank. High confidence reactions (*H*): The top 50 percentile of reactions with positive RIS values are included in the *H* set. These reactions are considered to have high confidence and are expected to be more relevant in the context-specific model. Medium confidence reactions (*M*): All other reactions with positive RIS values, excluding those already included in the *H* set, are included in the *M* set. These reactions are considered to have medium confidence and may have a lower impact or importance compared to the high confidence reactions. Reactions with no expression data (NER): Reactions that have no associated expression data are not included in either the *M* or *H* set.

By defining the *M* and *H* sets based on the positive RIS values and percentile ranking, the MBA method aims to prioritize reactions with higher expression levels and assign them different levels of confidence in terms of their relevance in the context-specific model.

## Conclusions

The proposed Localgini algorithm for thresholding gene expression data has shown promising results in building context-specific metabolic models. Localgini provides gene-specific thresholds for model construction By leveraging the Gini coefficient to capture the inequality in the dispersion of gene expression values. In this study, Localgini outperformed other thresholding methods, such as LocalT2 and StanDep, in terms of recovering the most number of housekeeping pathways and improving consensus among models. One of the key advantages of Localgini is its data-driven approach, which utilizes transcriptomics data to derive the thresholds. By considering the unique expression patterns of genes, Localgini provides a gene-specific thresholding strategy. This approach enhances the accuracy of the resulting context-specific models. Furthermore, Localgini contributes to reducing the contribution of MeMs in context-specific models by extracting more consistent reactions. This helps to streamline the model extraction process and prioritize reactions that are more consistently active across different contexts.

In summary, the Localgini algorithm provides a valuable tool for building context-specific metabolic models by leveraging gene-specific thresholds derived from the Gini coefficient. Its data-driven approach and ability to improve consensus among models make it a promising method for future research in the field of metabolic modeling.

## Supporting information

**S1 Text. Supporting text containing supplementary methods and results. S1 Fig. calculation of Gini coefficient from Lorenz curve**.

**S2 Fig. Extraction of context-specific models using Localgini thresholding method**. Complete pipeline demonstrating usage of Localgini threshold to extract context-specific models.

**S3 Fig. Number of housekeeping reactions rectified in core reaction list by each of the thresholding methods in the HPA dataset. LocalGini recovers the most number of housekeeping reactions in 31 tissues**

**S4 Fig. Localgini derived models are more self-consistent compared to other thresholding methods** Violin plots indicating the fraction of reactions added by MeMs for the given reaction importance by different thresholding methods in the HPA dataset.

**S5 Fig. Fraction of housekeeping reactions seeded in context-specific models built using different model extraction methods (MeM) and with three thresholding methods: Localgini, LocalT2, and StanDep in cancer cell-lines data**

**S6 Fig. Fraction of housekeeping reactions seeded in context-specific models built using different model extraction methods (MeM) and with three thresholding methods: Localgini, LocalT2, and StanDep in HPA data**

**S7 Fig. Localgini shows comparable performance in gene-essentiality predictions with other thresholding methods**

**S8 Fig. Localgini reduces false negatives in the core reaction list of the HPA dataset**. Pathways known to occur in the tissues are obtained from a manually curated resource and compared against the core reaction list obtained from different thresholding methods. Pathway enrichment analysis (hyper-geometric test) is done on the core reaction list for each of the tissue-pathway pairs. Different colours indicate that the core reaction list for the tissue is enriched with the reactions from the corresponding pathway (*p*-value*<* 0.05). The black color indicates none of the core reactions shows enrichment of the corresponding pathway.

**S9 Fig. Mean number of core reactions recovered by distinct thresholding methods in two different datasets used**.

**S10 Fig. Histogram of Jaccard similarity of models from same**

**cell-line/transcriptomics across different model extraction methods using Localgini, LocalT2, and StanDep thresholding approaches in cancer cell-lines data**

**S11 Fig. Histogram of Jaccard similarity of models from same**

**cell-line/transcriptomics across 3 different model extraction methods (FASTCORE, MBA, mCADRE) using Localgini, LocalT2, and StanDep thresholding approaches in cancer cell-lines data**

**S12 Fig. Histogram of Jaccard similarity of models from same**

**cell-line/transcriptomics across different model extraction methods using Localgini, LocalT2, and StanDep thresholding approaches in HPA data**.

**S13 Fig. Histogram of Jaccard similarity of models from same cell-line/transcriptomics across 3 different model extraction methods (FASTCORE, MBA, mCADRE) using Localgini, LocalT2, and StanDep thresholding approaches in HPA data**

**S14 Fig. Localgini recovers the most number of cancer hall-mark genes**

## Acknowledgments

We acknowledge the Centre for integrative biology and systems medicine (IBSE) for providing a high-performance cluster facility.

