## Supplemental text 1 for "Localgini: A method for harnessing inequality in gene expression to improve the quality of context-specific models"

### S1 Gini coefficient: An inequality metric to derive thresholds for context-specific model extraction

Efforts to enhance the accuracy of Genome-Scale Models (GSM) in capturing specific biological contexts necessitate the use of context-specific information as input. Recent advancements in high-throughput data generation, such as transcriptomics, have yielded valuable context-specific information for numerous human cells, encompassing various tissues, cell lines, and disease states. To construct context-specific models, Model extraction Methods (MeMs) rely on transforming this dataset into reaction importance scores. The mapping between gene expression data and reaction importance scores aims to assign higher importance to genes exhibiting consistent expression levels, such as housekeeping genes, across different tissues, while also considering context-specific reactions.

The Gini coefficient (GC) or Gini index [4] is a non-parametric measure widely used in economics to quantify the inequality or equality in the distribution of income among individuals in a given country or locality. Similarly, it can be utilized to assess the unequal distribution of any property among a set of samples [19, 25, 21, 22]. In the context of gene expression levels, the GC serves as a valuable tool for analyzing the distribution across different contexts. The GC is mathematically calculated and visualized using the Lorenz curve [11], where the cumulative fraction of samples/contexts are sorted in increasing order of their expression values on the x-axis, and the corresponding cumulative fraction of gene expression levels is plotted on the y-axis (see Figure (S1)). The GC is computed as the fraction of the area enclosed by the Lorenz curve and the line of equality. Theoretically, GC ranges from 0 (indicating perfect equality) to 1 (indicating perfect inequality) for non-negative gene expression values. In previous studies, the GC has been successfully applied to various RNA-seq datasets, enabling the identification of rare cell types [9], the assessment of heterogeneous expression patterns of transporter proteins, the identification of housekeeping genes [12], and the selection of reference genes for expression profiling [26]. The Localgini approach leverages the GC to derive gene-specific thresholds from RNA-seq data, facilitating the construction of context-specific models.

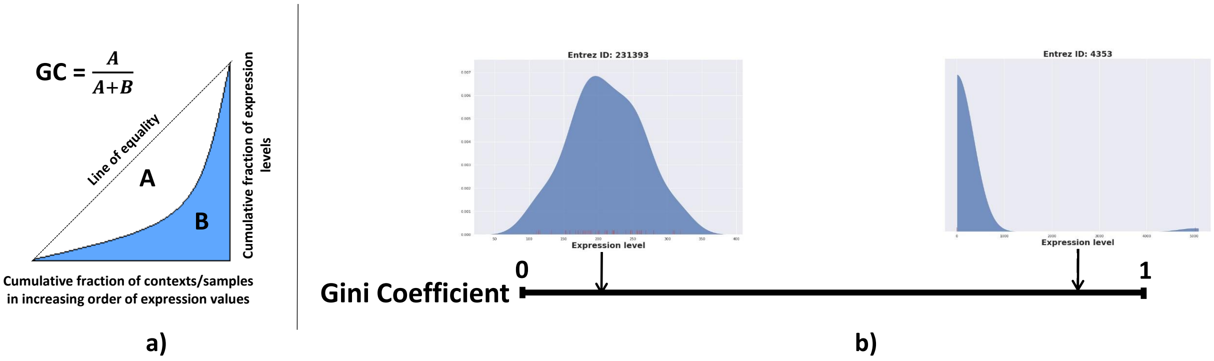

Figure S1: a) calculation of gini coefficient from Lorenz curve b) In Recon2.2, the GC values of genes ranged from 0.137 to 0.98

### S2 Supplementary methods

#### S2.1 Data processing

The Recon2.2 model incorporates Gene-Protein-Reaction (GPR) rules that provide information about the association between genes, proteins, and reactions within the model. These rules specify the isoforms or subunits of an enzyme required to catalyze a particular reaction. In GPR rules, ‘isoforms’, are typically indicated by an ‘OR’ relation between genes, implying that any one of the genes can be responsible for the enzymatic activity. On the other hand, the ‘subunits’ of a multimeric enzyme are indicated by an ‘AND’ relation between genes, signifying that all the genes must be present for the enzyme to be functional. In the context of gene expression data, thresholds are typically defined at the gene, enzyme, or reaction level. These thresholds are used to determine the activation or inactivation status of genes, enzymes, or reactions based on their expression levels.

##### S2.1.1 Gene level

In the LocalT2 method, the threshold is applied at the gene expression level. The gene importance or core gene set is determined based on whether the gene’s expression value exceeds the specified threshold. To map this gene importance/core gene set to reactions in the model, the associated Gene-Protein-Reaction (GPR) rules are utilized. For GPR rules with an ‘AND’ relation between genes, the expression mapping is done by taking the minimum expression value among the genes involved in the rule. This implies that all the genes in the ‘AND’ rule must have expression values above the threshold for the reaction to be considered as part of the core reaction set. Conversely, for GPR rules with an ‘OR’ relation between genes, the expression mapping is done by taking the maximum expression value among the genes involved in the rule. This means that if any of the genes in the ‘OR’ rule have expression values above the threshold, the reaction is considered part of the core reaction set. By applying these expression mappings based on the GPR rules, the reaction importance or core reaction set is obtained for the LocalT2 method [16].

##### S2.1.2 Enzyme level

In the StanDep method, the thresholding is performed at the enzyme level. The gene expression data is used to calculate the enzyme expression values, which are then subjected to thresholding [10]. To map the gene expression data to enzyme expression values, the ‘AND’ rule in the GPR is resolved by taking the minimum expression value among the genes involved in the rule. This means that all genes in the ‘AND’ rule must have expression values above the threshold for the corresponding enzyme to be considered active. Unlike in the case of ‘AND’ rules, the ‘OR’ rule is not resolved at the enzyme level. Instead, the isoforms associated with the enzyme are included in the enzyme expression dataset, regardless of their individual gene expression values. Once the enzyme expression values are obtained, the threshold is applied to identify the enzyme importance or core enzyme set. The enzyme importance/core enzyme set is then mapped to the reactions using the GPR rules to obtain the corresponding reaction importance or core reaction set.

##### S2.1.3 Reaction level

In the process of mapping gene expression values to reactions using GPR rules, the ‘AND’ rule is resolved by taking the minimum expression value among the genes involved in the rule. On the other hand, the ‘OR’ rule can be resolved in different ways. One approach is to take the maximum expression value among

the genes involved in the 'OR' rule. This means that if any of the genes in the 'OR' rule have expression values above the threshold, the corresponding reaction will be considered active. Alternatively, the 'OR' rule can be resolved by taking the sum of the expression values of the genes involved. In this case, if the sum of expression values exceeds the threshold, the reaction will be considered active. Once the reaction expression data is obtained by mapping gene expression values using the GPR rules, thresholds are provided to this data to define the reaction importance or active reaction set. Reactions with expression values above the threshold are considered important or active, while reactions with expression values below the threshold are considered to be inactive.
In a previous study, ([17]), it was demonstrated that gene-level thresholding outperformed reaction-level thresholding in terms of performance. Based on this finding, the Localgini method was specifically implemented at the gene level for this study. However, it's worth noting that the Localgini algorithm can be adapted and implemented at different levels, including gene, enzyme, and reaction levels. The code '`buildContextmodels.m`' can be used to apply thresholding at different levels and get the corresponding models.

#### S2.2 Thresholding methods

Thresholding is a crucial step in extracting context-specific models from generic genome-scale models, such as Recon X or iHsa. Model extraction methods rely on pre-defined sets of active reactions to construct these context-specific models. To determine which reactions should be considered active, thresholds are applied to the expression data, above which the reactions or genes are considered active. However, the selection of these thresholds is often done arbitrarily without a systematic approach. In this study, we introduce a novel thresholding algorithm called Localgini, which utilizes the Gini coefficient of genes to derive context-specific thresholds. The Gini coefficient is a measure of inequality/equality in the distribution of gene expression levels across different contexts. By analyzing the distribution patterns of gene expression, Localgini calculates gene-specific thresholds that reflect the variability and inequality in gene expression across contexts. The overall workflow for building context-specific models from Recon2.2 and RNAseq data is illustrated in Figure (S2). This workflow incorporates the Localgini algorithm to determine the active reactions based on the derived thresholds.

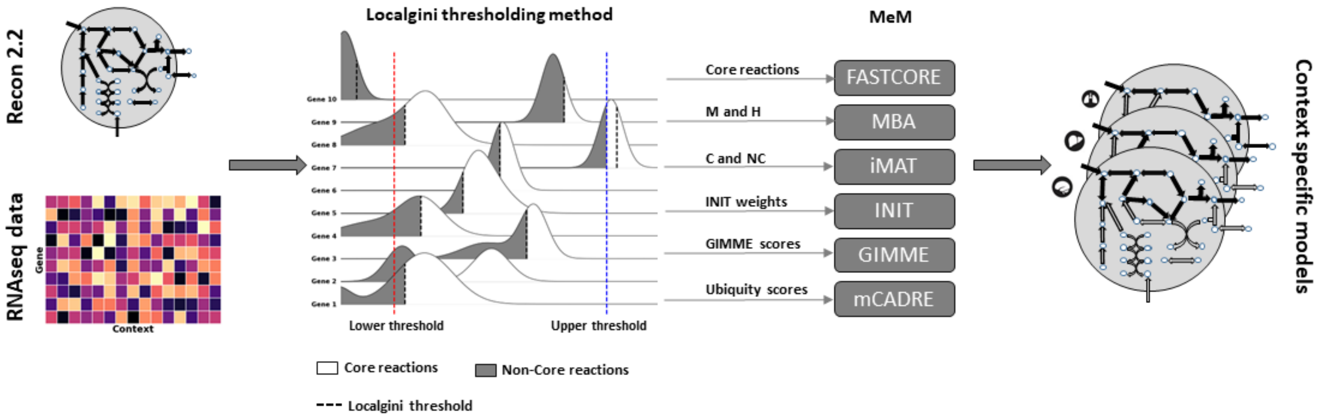

Figure S2: Extraction of context-specific models using Localgini thresholding method

###### S2.2.1 Global threshold

Traditionally, a common approach to define active reactions or genes from gene-expression data is to apply a single threshold to the entire dataset. This global thresholding method categorizes genes as active if their expression values exceed the threshold, while genes with expression values below the threshold are considered inactive. Consequently, reactions associated with the active genes are labeled as active reactions. However, this global approach assumes a uniform expression pattern for all genes, which oversimplifies the complexity of gene expression regulation across different contexts. In contrast, methods like LocalT2 and StanDep have been proposed as alternatives to the global approach and have demonstrated improved performance in previous studies ([10]). These methods take into account the context-specific variations in gene expression and consider the local neighborhood of each gene to determine its activity.

###### S2.2.2 LocalT2 threshold

LocalT2 is a gene-specific thresholding method that assigns a unique threshold to each gene based on its mean expression value across contexts. This gene-level threshold is determined as the average expression value of the gene across all contexts. Additionally, two global thresholds, namely the lower and upper thresholds, are defined to selectively include highly over-expressed reactions and exclude under-expressed reactions. In the study by [16], these global thresholds are set based on the $25^{\mathtt{th}}$ and $75^{\mathtt{th}}$ percentiles of the expression values, respectively.

By utilizing gene-specific thresholds in combination with global thresholds, LocalT2 takes into account both the individual gene expression patterns and the overall distribution of expression values across all genes. This approach allows for the identification of genes that exhibit exceptionally high or low expression levels relative to the average, thereby capturing the full range of gene expression dynamics in context-specific models.

###### S2.2.3 StanDep threshold

In the study by [10], a novel thresholding algorithm called StanDep was developed to define thresholds for enzymes based on their expression patterns. Unlike the global thresholding method that provides a single threshold for all genes or the local methods that assign a gene-specific threshold, StanDep takes an intermediate approach by providing cluster-specific thresholds. The StanDep algorithm follows a series of steps to determine the cluster-specific thresholds:

The number of enzyme clusters is chosen such that the addition of one more cluster does not significantly alter the core reaction list, ensuring a high degree of similarity (more than 90

- Gene expression values are mapped to the enzyme expression values and transformed using a logarithmic function ($\mathtt{log}_{10}$).
- The expression values of each enzyme are then binned into predefined bins, resulting in a vector of counts for each bin.
- Hierarchical clustering is performed on these count vectors to group enzymes based on their expression patterns.
- For each enzyme cluster, a cluster-specific threshold is calculated using Equation (S1).

- The number of enzyme clusters is chosen such that the addition of one more cluster does not significantly alter the core reaction list, ensuring a high degree of similarity (> 90% similarity).

Cluster-specific threshold is calculated by,

$$\begin{aligned}
\Theta_c &= \frac{(\theta_c - \min(\theta_c))100}{\max(\theta_c - \min(\theta_c))}, \\
\theta_c &= f(\sigma_c) + g(\mu_c), \\
f(\sigma_c) &= \frac{(\sigma_c - \Delta)}{\max(\sigma_c - \Delta)} \\
g(\mu_c) &= (\mu_c - M) \qquad \text{(S1)}
\end{aligned}$$

where

$\Theta_c$ = Processed threshold value for the cluster $c$; $\theta_c$ = Raw threshold value for the cluster $c$; $\sigma_c$ = Standard deviation of the cluster $c$; $\Delta$ = Standard deviation of the dataset ; $\mu_c$ = Mean of the cluster $c$ ; $M$ = Mean of the dataset

The bin edges used for datasets are as follows: for NCI60 cancer cell lines dataset,[-3, -2, -1, 0, 1, 2, 2.5, 3, 4], and for HPA datset, [-3, -2, -1, 0, 1, 2, 2.5, 3, 4, 5]. For hierarchical clustering, the Euclidean distance metric and complete linkage metric are used for both datasets. The total number of clusters obtained were 26 and 19 for NCI60 dataset and HPA dataset, respectively.

#### S2.3 Implementation of Localgini with MeMs

##### S2.3.1 FASTCORE

In the FASTCORE algorithm developed by [23], the core reactions that are deemed to be present in a specific context are provided as input. The algorithm aims to identify the minimal set of reactions that can support these core reactions and includes them in the context-specific model. In the case of this study, the core reactions are defined based on the Localgini thresholds. Specifically, all reactions with expression values greater than the gene-specific thresholds derived from the Localgini algorithm are considered as core reactions. Additionally, the biomass function and ATP demand reactions are manually added to the core reaction set, as they are essential for cellular function. However, certain reactions are excluded from the core reaction list based on specific criteria. Reactions that lack information on Gene-Protein-Reaction (GPR) associations in the Recon2.2 model are not included. Furthermore, reactions for which no gene expression data is available, referred to as Non-Expressed Reactions (NER), are also excluded.

##### S2.3.2 iMAT

The iMAT algorithm, introduced by [27], requires two sets of reactions, namely core ($C$) and non-core ($NC$), to be provided as input. The algorithm aims to maximize the inclusion of core reactions while excluding non-core reactions from the context-specific model. In this study, the core reaction set ($C$) is defined based on the Localgini thresholds. Reactions with expression values greater than the gene-specific thresholds derived from the Localgini algorithm are included in the core set. Additionally, the biomass function and ATP demand reactions are manually added to the core set, as they are essential for cellular function. On the other hand, the non-core reaction set ($\mathsf{NC}$) consists of reactions with expression values lesser than the Localgini thresholds. It's important to note that reactions corresponding to NER, for which no gene expression data is available, are not included in either the core or non-core sets.

##### S2.3.3 INIT

The INIT algorithm, as described by [1], requires the provision of reaction weights to determine the importance of each reaction. The algorithm aims to maximize the inclusion of positively weighted reactions while minimizing or removing negatively weighted reactions from the context-specific model. In this study, the reaction importance scores (RIS), as explained in Section 2.4.2 of the main text, are used as the weights for INIT. The maximum RIS value among the reactions is assigned as the weight for the biomass function and ATP demand reactions, indicating their high importance. For NER, which are reactions with no available gene expression data, a weight of 0 is assigned. This reflects the negligible contribution of NER for model extraction.

##### S2.3.4 MBA

The MBA algorithm, proposed by [8], requires the input of two sets of reactions: medium confidence reactions ($\mathsf{M}$) and high confidence reactions ($\mathsf{H}$). In the context-specific model, high-confidence reactions are always included, while medium-confidence reactions are included only if a specific trade-off criterion is met in terms of excluding non-core reactions. All other reactions are included if they support the high-confidence reactions. The top 50 percentile of reactions with positive RIS is included in the set $\mathsf{H}$. This means that reactions with RIS values above a certain threshold are considered high-confidence reactions. Additionally, the biomass function and ATP demand reactions are manually included in the set $\mathsf{H}$ to ensure their presence in the model. All other reactions with positive RIS values, excluding the high-confidence reactions, are included in the set $\mathsf{M}$. NER, which are reactions with no available gene expression data, are not included in either set $\mathsf{H}$ or set $\mathsf{M}$. The parameter $\epsilon$ is used to determine the trade-off between the inclusion of medium confidence reactions ($\mathsf{M}$) and the exclusion of non-core reactions. In this study, $\epsilon$ is set to 0.5, indicating that the inclusion of medium-confidence reactions is considered only if it does not compromise the exclusion of non-core reactions.

##### S2.3.5 GIMME

The GIMME (Gene Inactivity Moderated by Metabolism and Expression) algorithm, introduced by [3], requires two inputs: gimme_scores and a threshold value. The final context-specific model is constructed by including reactions with gimme_scores greater than the threshold value. All other reactions are pruned based on their scores and their connectivity to the biomass function.

Gimme_scores are calculated for each reaction in the model. These scores are similar to the reaction importance scores (RIS) mentioned earlier, with the exception that Non-Expressed Reactions (NER) are assigned a score of $-\epsilon$, where $\epsilon$ is a small positive number. The maximum RIS value is assigned as the gimme_score for the biomass function and ATP demand reactions. The input threshold is given as 0.

##### S2.3.6 mCADRE

To apply the mCADRE algorithm, as described by [24], we need to provide three inputs: ubiquity scores ($U$), confidence scores ($C$), and a threshold value. Ubiquity scores ($U$) are calculated for each reaction in the model. In this case, the ubiquity scores are derived from the reaction importance scores (RIS) mentioned earlier. Reactions with positive RIS, as well as the biomass function and ATP demand reactions, are assigned a ubiquity score of 1. Non-Expressed Reactions (NER) are given a ubiquity score of $1-\epsilon$, where $\epsilon$ is a small positive number. For other reactions, the RIS values are min-max scaled to range from 0 to $1-\epsilon$, ensuring that the ubiquity scores lie within this range. The confidence scores ($C$) are assigned to each reaction, representing the prior confidence in the presence of the reaction in the context. In this case, as there is no prior information available for the inclusion of a reaction, the confidence scores for all reactions are set to 0. The threshold value is given as 1. Reactions with ubiquity scores greater than or equal to the threshold are considered core reactions and are included in the final model.

#### S2.4 Extracting housekeeping reactions in Recon2.2

To identify housekeeping reactions, a list of 3804 genes that are uniformly expressed across tissue samples was obtained from [5]. This gene list was then converted to Entrez ID format using the Biomart API [20]. Out of these genes, 318 housekeeping genes were found to be present in the flux-consistent model of Recon2.2. To extract housekeeping reactions from Recon2.2, the rxnGeneMat matrix was used. The rxnGeneMat is a binary matrix where the rows represent reactions and the columns represent genes. A value of 1 indicates the existence of a relationship between the reaction and the gene, while a value of 0 indicates no relationship. In this case, all rows with non-zero elements in the columns corresponding to the housekeeping genes were considered as housekeeping reactions. As a result, a total of 929 housekeeping reactions were identified in Recon2.2.

#### S2.5 Fractional contribution of reactions by MeMs in the final model

Self-consistency is the property of the thresholding method, which checks the contribution of the reactions by the MeM and the thresholding method in the final model. The fractional contribution of a Model extraction Method (MeM), denoted as $f_{MeM,M}$, in the final model M for a given thresholding method can be calculated as follows:

$$f_{MeM,M} = \frac{card(M_{rxn} \setminus T_{rxn})}{card(M_{rxn})} \tag{S2}$$

where

$M_{rxn}$ = Set of reactions in M;
$T_{rxn}$ = Set of core reactions as provided by the thresholding method;
$card(.)$ = Cardinality of the set.

This calculation measures the proportion of reactions in the final model that were extracted by the specific MeM. It provides an indication of the contribution of the MeM in determining the reaction set of the model when combined with the thresholding method.

#### S2.6 Essential genes prediction using fastSL

Flux Balance Analysis (FBA) based simulations on a metabolic model can predict the genes necessary for the growth of the cell [13]. FBA formulation to predict the essential genes in a metabolic model is given below:

$$\begin{aligned} \text{max.} \quad & \nu_{bio} \\ \text{subject to} \quad & \sum_j s_{ij}\nu_j = 0, \quad \forall i \in M, \forall j \in R \\ & LB_j \leqslant \nu_j \leqslant UB_j, \quad \forall j \in R \\ & \nu_d = 0, \quad d \in D \subset R \end{aligned} \tag{S3}$$

where

$\nu_{bio}$ = Flux through the biomass reaction;
$s_{ij}$ = Stoichiometric coefficient of $i^{th}$ metabolite in $j^{th}$ reaction;
$\nu_j$ = Flux through reaction $j$;
$M$ = Set of all the metabolites in the model;
$R$ = Set of all reactions in the model;
$LB_j$ = Lower bound of flux through $j^{th}$ reaction;
$UB_j$ = Upper bound of flux through $j^{th}$ reaction;
$D$ = Set of all the reactions that are to be deleted after a gene removal.

To predict whether a gene $G$ is essential for biomass growth, maximum biomass growth flux is predicted by solving the linear program S3 for the wild type model ($D = \emptyset$) and the mutant model ($D$ = reactions that are dependent on Gene $G$). Standard formulation iterates through all the genes, hence corresponding reactions in the model, and checks the impairment of growth before and after removal of the gene to predict its essentiality. FastSl [15] prunes the search space by providing a set of reactions $J_{nz}$ that has the essential genes as its subset. To identify $J_{nz}$ fastSL computes minimum $l_1$-norm solution of the following optimization problem.

$$\begin{aligned} \text{min.} \quad & \sum_j |\nu_j| \\ \text{subject to} \quad & \sum_j s_{ij}\nu_j = 0, \quad \forall i \in M, \forall j \in R \\ & LB_j \leqslant \nu_j \leqslant UB_j, \quad \forall j \in R \\ & \nu_{bio} = \nu_{bio,WT} \end{aligned} \tag{S4}$$

Reactions with non-zero flux in the steady-state flux distribution are taken in set $J_{nz}$

#### S2.7 Metabolic task

Metabolic capability is an important aspect of evaluating a metabolic model. It assesses the model's ability to perform specific metabolic tasks, such as producing certain products from defined input substrates [2] [14]. For example, ATP generation from glucose requires glucose, oxygen, phosphate, ADP, and hydrogen as input and output should be ATP, $H_2O$, and $CO_2$. The evaluation is typically done by formulating a linear programming problem, where the only exchange reactions with non-zero flux bounds are the specified input substrates and output products. To evaluate the metabolic capability of a model, a curated list of metabolic tasks is used. These tasks cover various aspects of cellular metabolism, including energy generation, carbohydrate metabolism, vitamin and cofactor metabolism, lipid metabolism, nucleotide metabolism, amino acid metabolism, and glycan metabolism. This list of metabolic tasks was manually curated by Richelle et al. [16] and consists of 210 distinct tasks. The COBRA toolbox [7], provides a function called *checkMetabolicTasks.m.* This function is used to test whether a given model passes each of the metabolic tasks defined in the curated list. It checks if the linear programming problem for each task is solvable, and if so, it marks the task as passed (assigned a value of 1), otherwise, it marks it as not passed (assigned a value of 0). By running the *checkMetabolicTasks.m* function for a set of models, a binary matrix can be created. Rows in the matrix represent different models, while columns represent the metabolic tasks. The elements of the matrix indicate whether a model passed a specific task (1) or not (0). This matrix can then be used for further analysis to identify the causes of variability in model performance and to compare the metabolic capabilities of different models.

#### S2.8 Principal component analysis on reaction content and metabolic functionalities

The evaluation of thresholding methods for reducing variance across models built from the same transcriptomics data involves analyzing the variation in model content, and metabolic functionality [6][18]. One approach to increase consensus among models is to assign higher importance scores to reactions or pathways known to occur in the specific context if such prior knowledge is available. To assess the ability of Localgini and other thresholding methods to reduce variance occurred due to different MeMs, Principal Component Analysis (PCA) is performed on the matrices representing the reaction content and metabolic functionality of the models. PCA is a statistical technique used to reduce the dimensionality of a dataset while retaining the most important information. In this evaluation, the Pearson correlation coefficient (PCC) is calculated between the principal components obtained from PCA and the categories of interest, including cell-line/transcriptomic data, MeMs, and cancer types. For example, for the category of 44 cancer cell-lines, each cell-line is assigned a unique random number from 1 to 44. The PCC is calculated between the principal component and the encoded cell-line category. This process is repeated 10,000 times to obtain a distribution of PCC values, and the maximum PCC value is squared, scaled to a percentage, and considered as the variance explained by the cell-lines in the corresponding principal component. The same approach is followed for MeMs and cancer types, where MeMs are encoded from 1 to 6 and cancer types are encoded from 1 to 9. By calculating the PCC between the principal components and the encoded categories, the variance explained by MeMs and cancer types can be determined.

Overall, this analysis allows for the quantification of the extent to which the thresholding methods, including Localgini, reduce variance across models built from the same transcriptomics data by examining the relationship between the principal components and the relevant categories of interest.

#### S2.9 Jaccard similarity between the models

Jaccard similarity (JC)/ Jaccard index between models built from the same cell-line/transcriptomics data is calculated for all three thresholding methods. To calculate the Jaccard similarity between the models M1 and M2, two binary vectors $R_{M1}$ and $R_{M2}$ of size $r \times 1$ are taken. $r$ is the number of reactions in Recon2.2. Each element in $R_{M1}$ and $R_{M2}$ refer to the presence (1) or absence (0) of the corresponding reaction in M1 and M2 respectively. JC between M1 and M2 is calculated using the below formula,

$$JC = \frac{R_{M1} \times R_{M2}^{T}}{\sum_{i=1}^{r}(R_{M1,i} + R_{M2,i}) - (R_{M1} \times R_{M2}^{T})} \tag{S5}$$

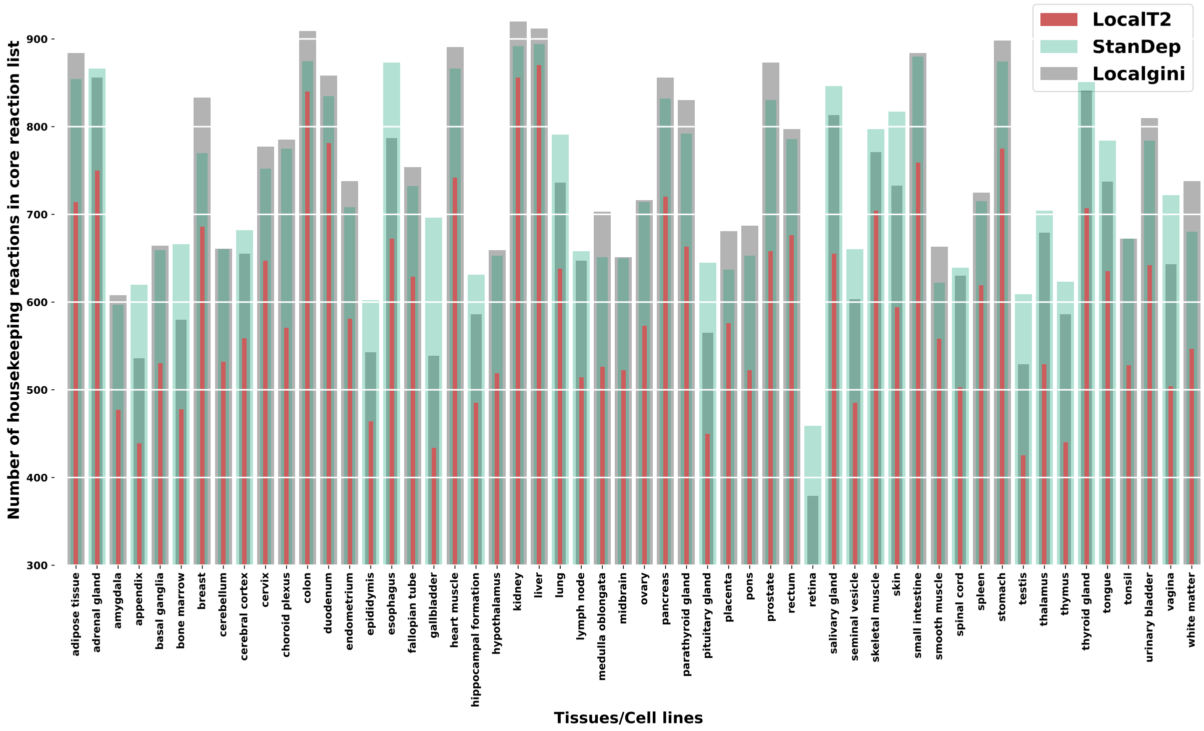

Figure S3: Number of housekeeping reactions rectified in core reaction list by each of the thresholding methods in the HPA dataset. LocalGini recovers the most number of housekeeping reactions in 31 tissues while StanDep recovers the most number of housekeeping reactions in 23 tissues.

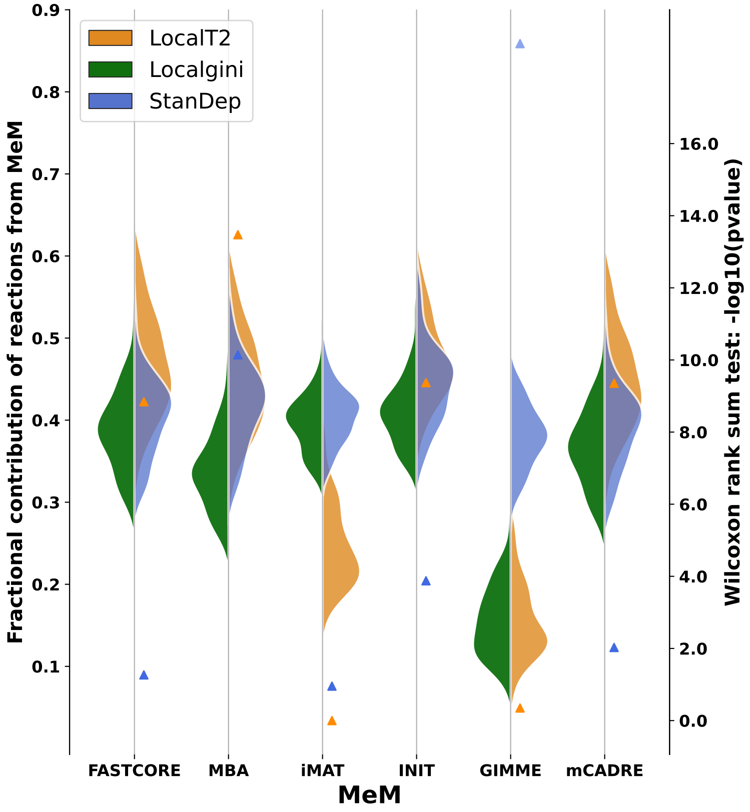

Figure S4: **Localgini derived models are more self-consistent compared to other thresholding methods** Left y-axis: Violin plots indicating the fraction of reactions added by MeMs for the given reaction importance by different thresholding methods in HPA dataset. Right y-axis: Triangles indicating the $-\log_{10}(p-value)$ of the left-tailed Wilcoxon rank sum test to compare the distribution of Localgini with LocalT2 (orange colour) and StanDep (blue colour)

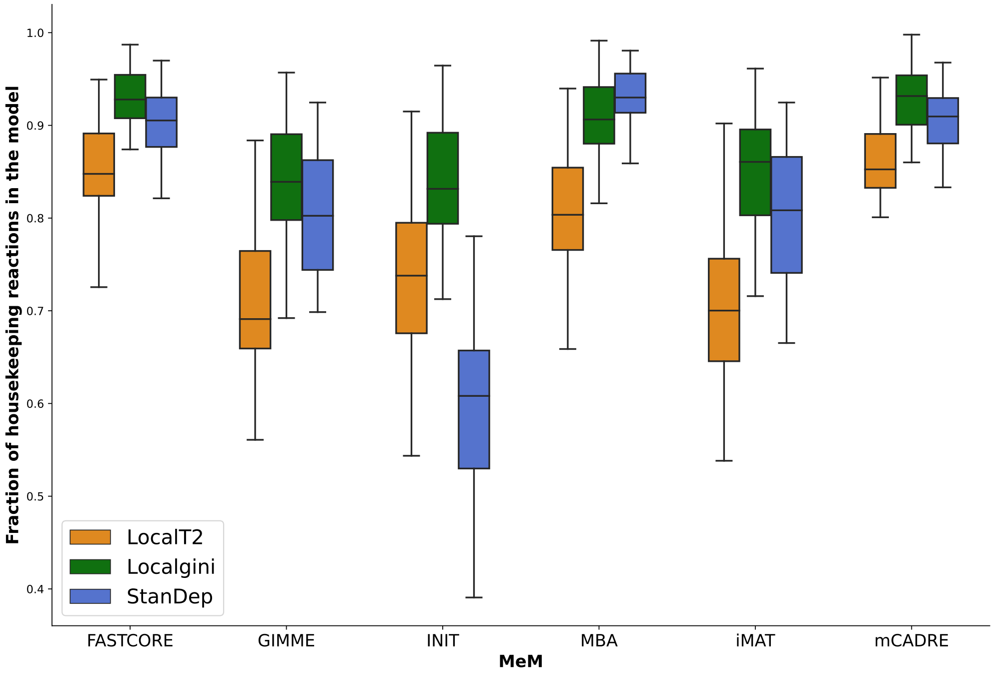

Figure S5: Fraction of housekeeping reactions seeded in context-specific models built using different model extraction methods (MeM) and with three thresholding methods: Localgini, LocalT2, and StanDep in cancer cell-lines data

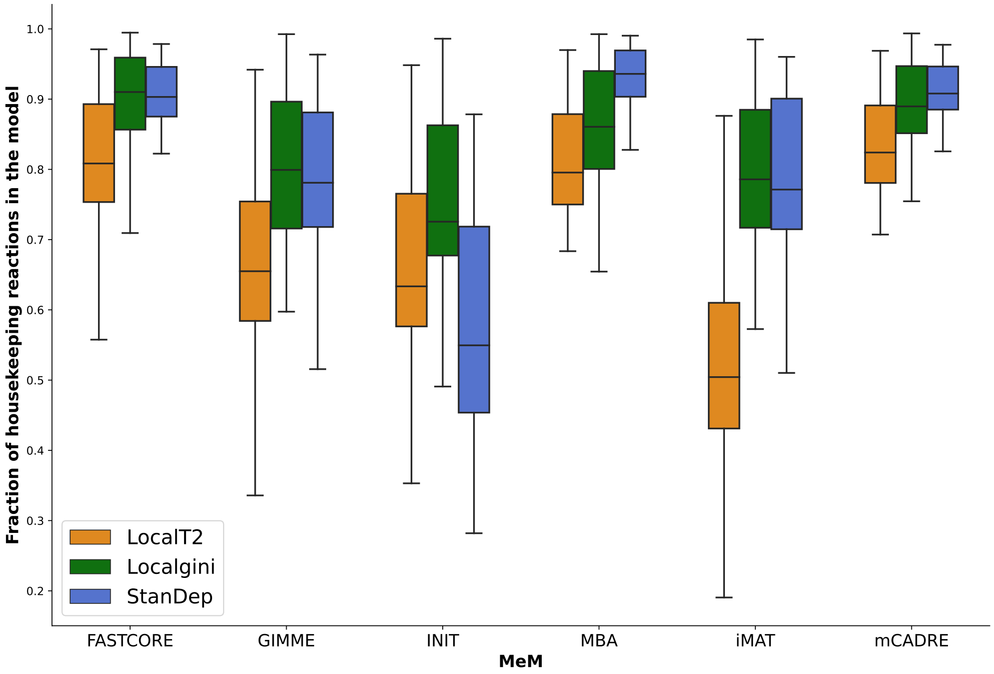

Figure S6: Fraction of housekeeping reactions seeded in context-specific models built using different model extraction methods (MeM) and with three thresholding methods: Localgini, LocalT2, and StanDep in HPA data

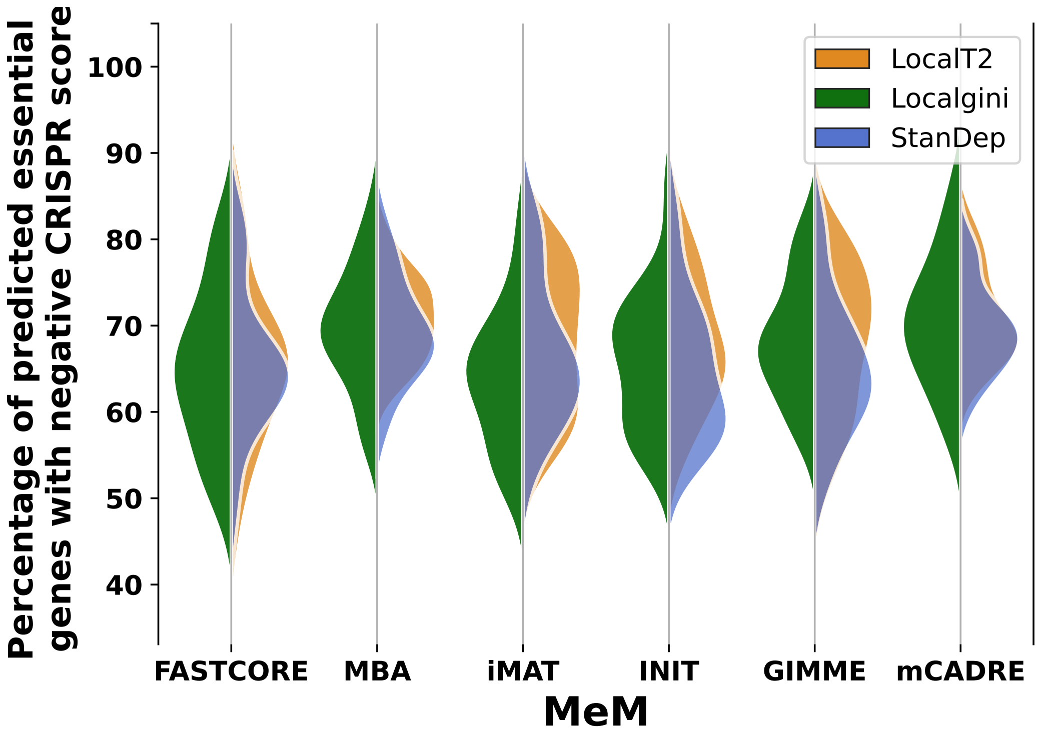

Figure S7: Localgini shows comparable performance in gene-essentiality predictions with other thresholding methods

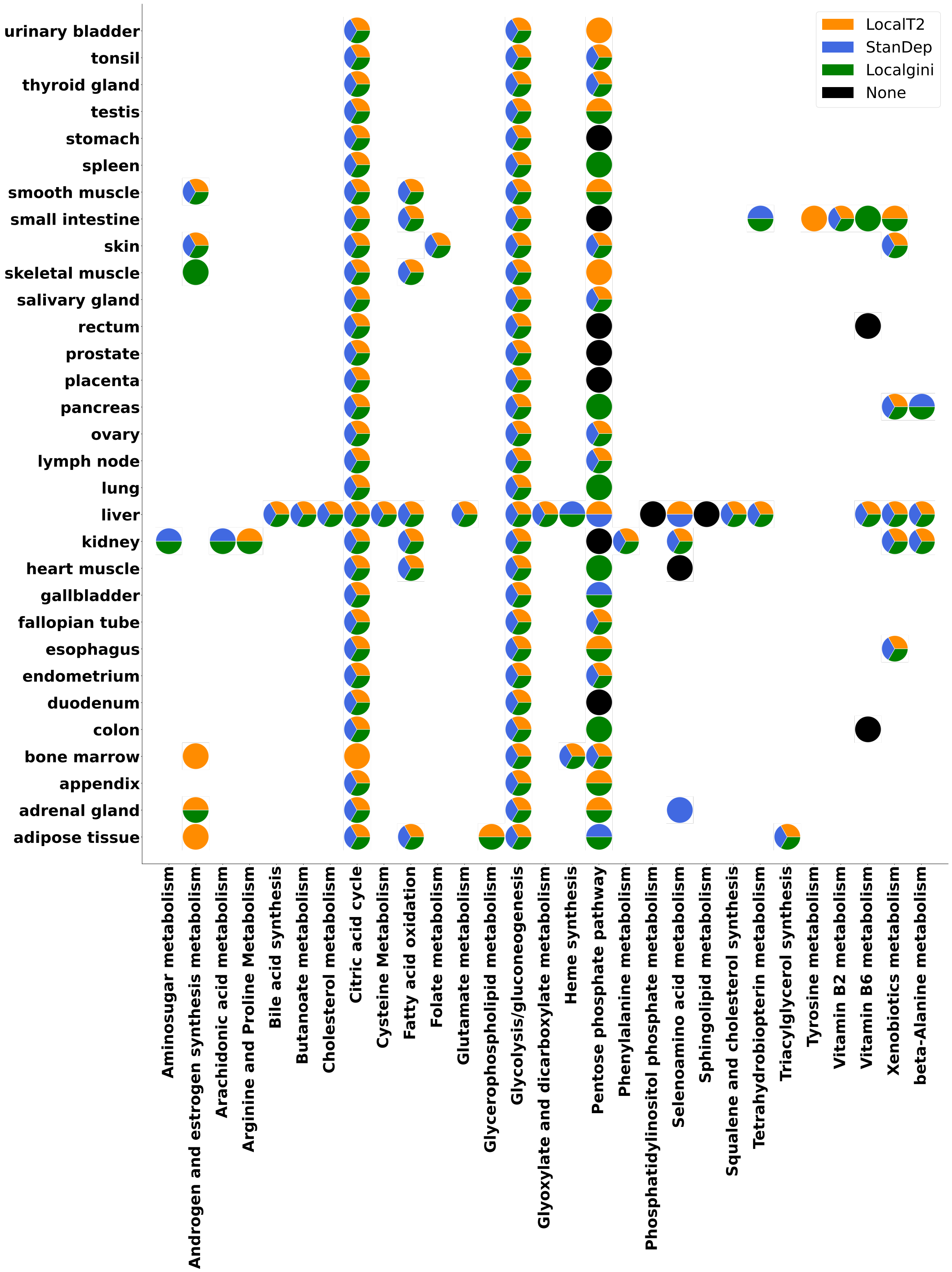

Figure S8: Pathways known to occur in the tissues are obtained from a manually curated resource([17]) and compared against the core reaction list obtained from different thresholding methods. Pathway enrichment analysis (hyper-geometric test) is done on the core reaction list for each of the tissue-pathway pairs. Different colours indicate that the core reaction list for the tissue is enriched with the reactions from corresponding pathway (p-value<0.05). Black color indicates none of the core reactions shows enrichment of the corresponding pathway

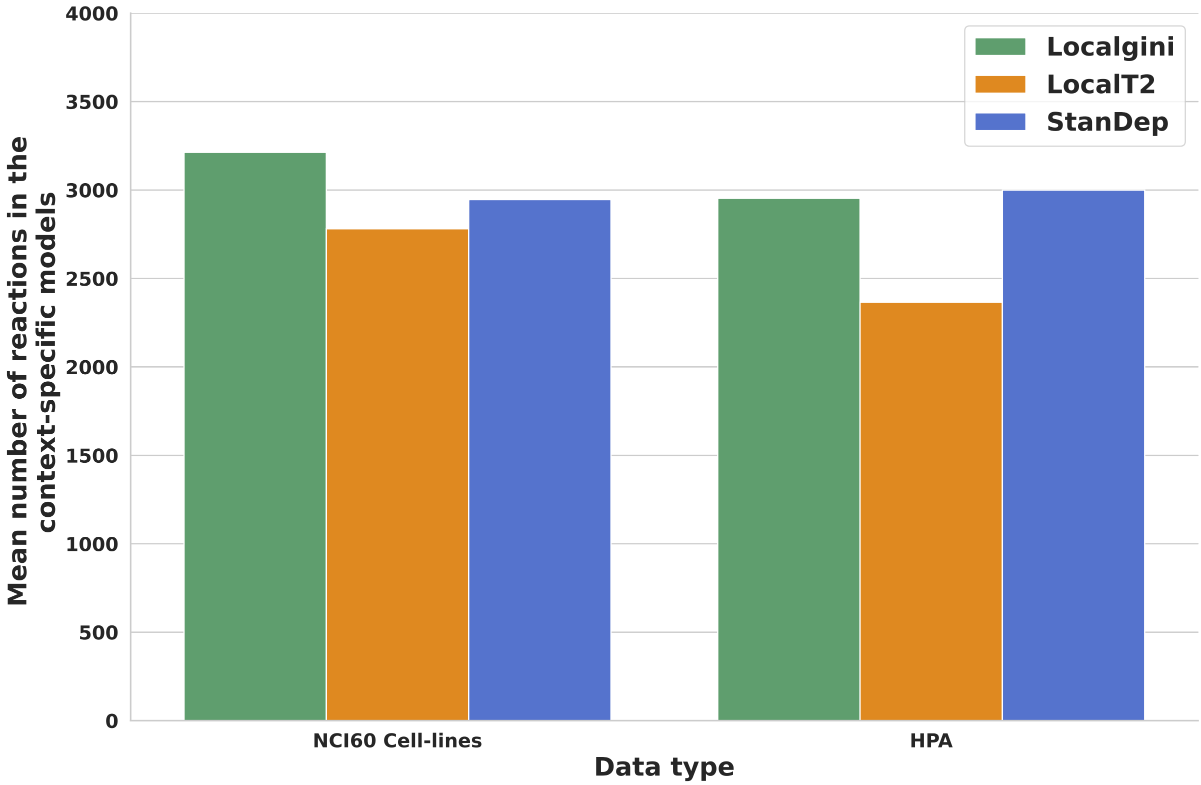

Figure S9: Mean number of reactions present in context-specific models built by distinct thresholding methods in two different datasets used

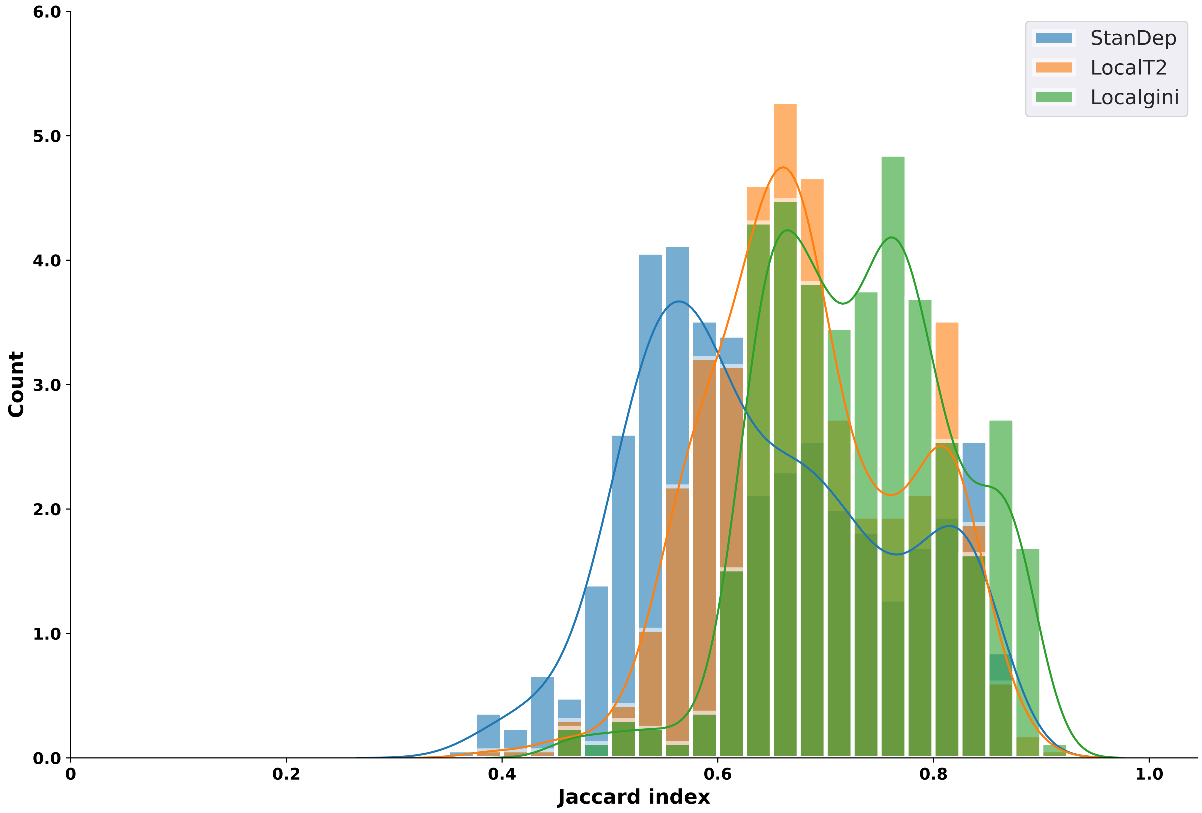

Figure S10: Histogram of Jaccard similarity of models from same cell-line/transcriptomics across different model extraction methods using Localgini, LocalT2, and StanDep thresholding approaches in cancer cell-lines data

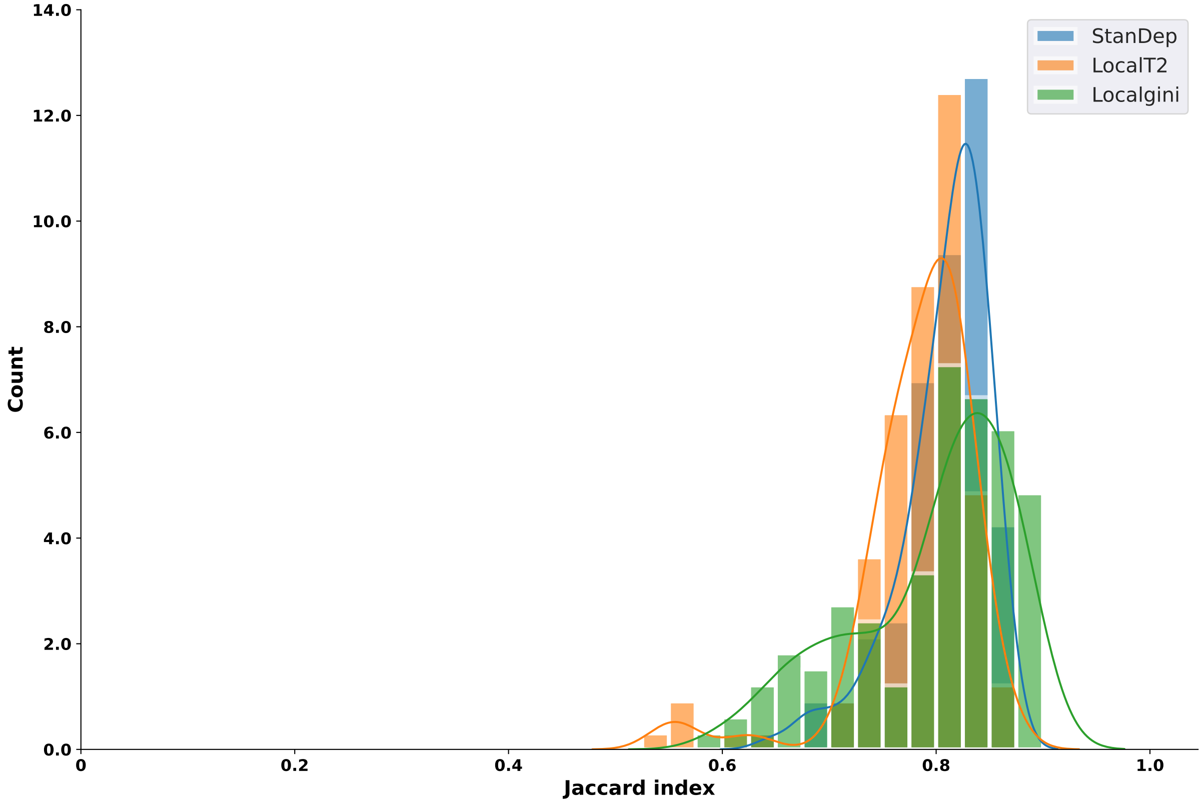

Figure S11: Histogram of Jaccard similarity of models from same cell-line/transcriptomics across 3 different model extraction methods (FASTCORE, MBA, mCADRE) using Localgini, LocalT2, and StanDep thresholding approaches in cancer cell-lines data

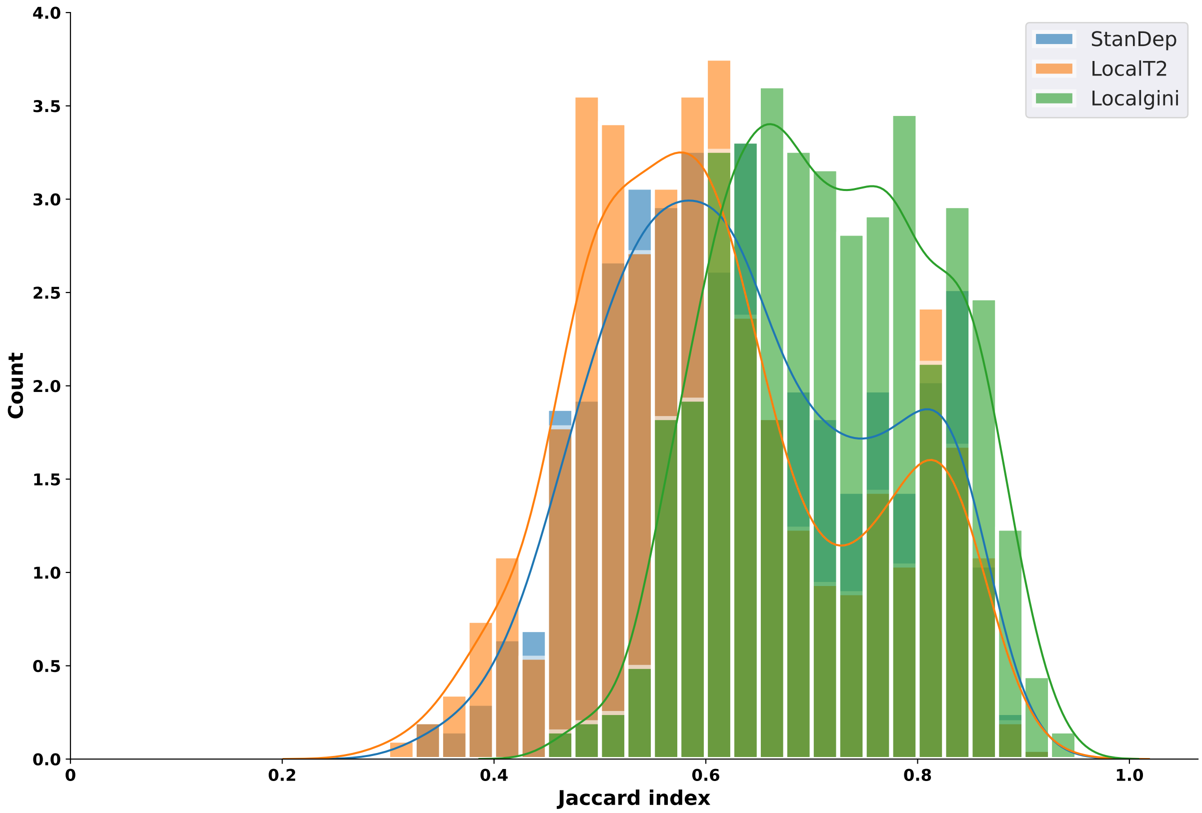

Figure S12: Histogram of Jaccard similarity of models from same cell-line/transcriptomics across different model extraction methods using Localgini, LocalT2, and StanDep thresholding approaches in HPA data

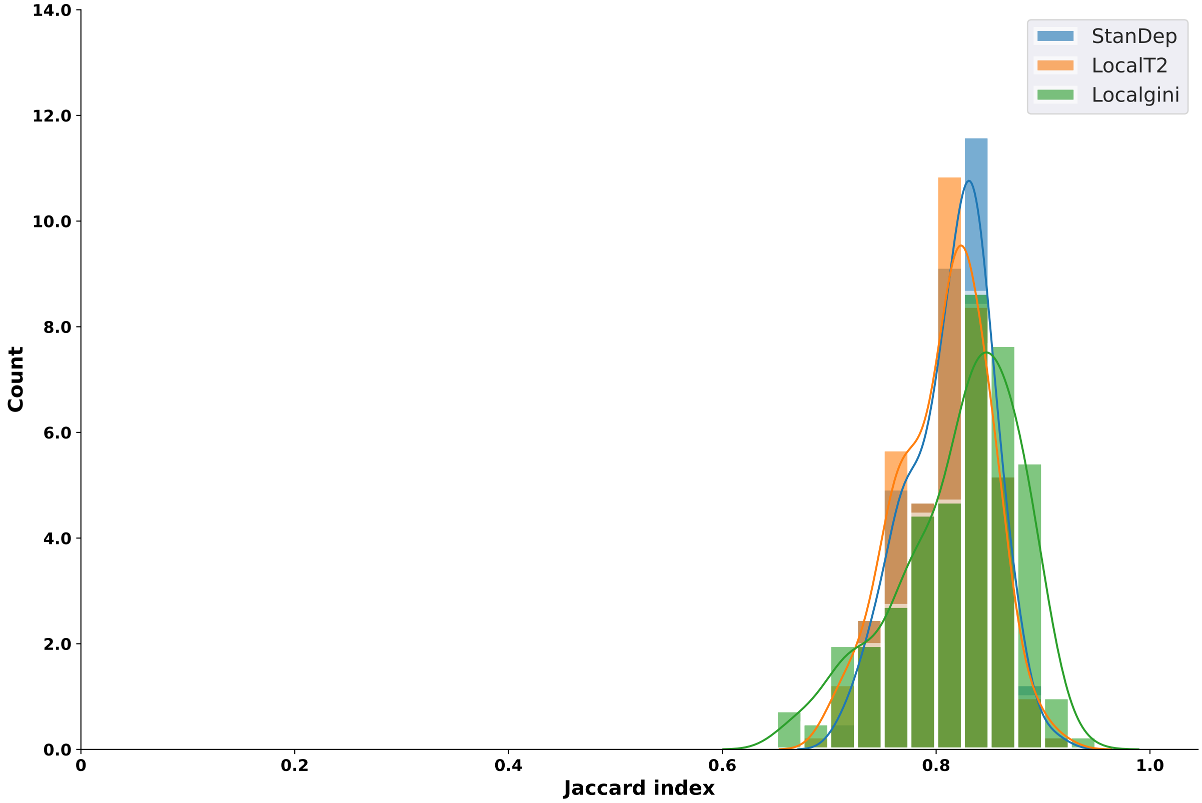

Figure S13: Histogram of Jaccard similarity of models from same cell-line/transcriptomics across 3 different model extraction methods (FASTCORE, MBA, mCADRE) using Localgini, LocalT2, and StanDep thresholding approaches in HPA data

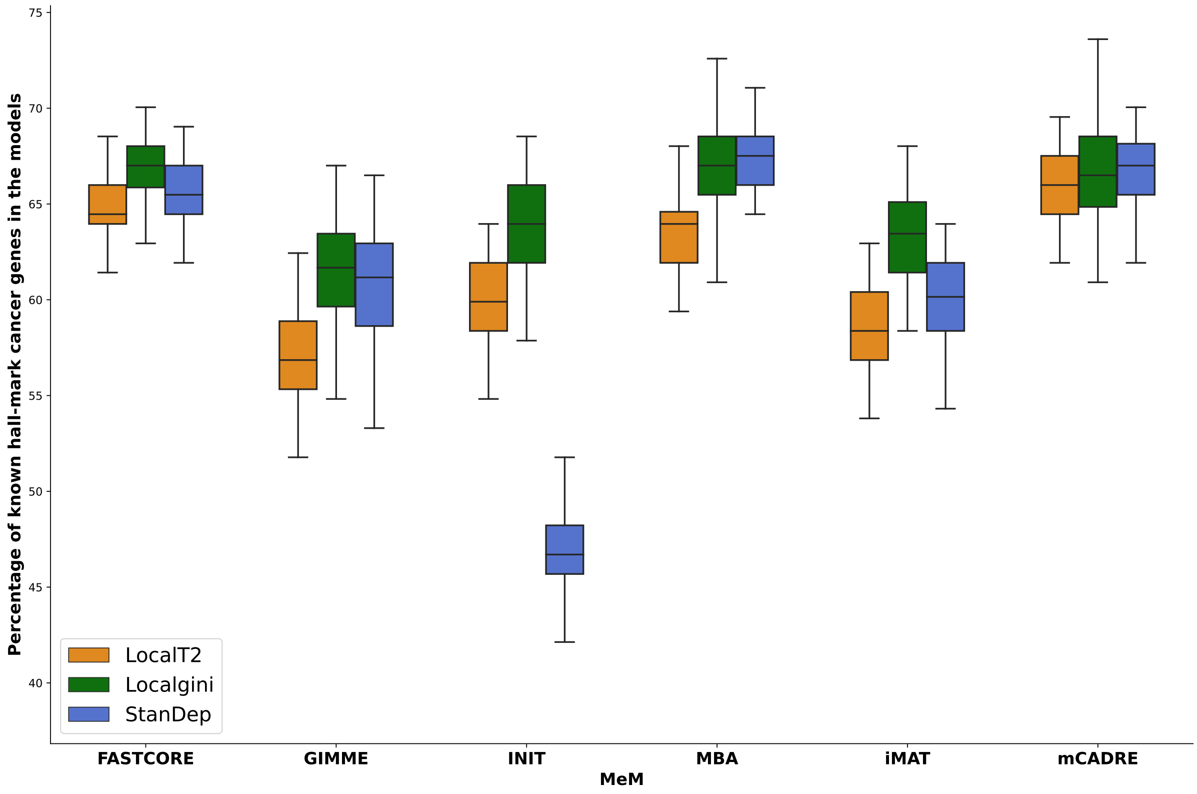

Figure S14: Localgini recovers most number of cancer hall-mark genes in most of the MeMs
